# S-acylation and tonoplast localization of the Geminivirus Rep-Interacting Kinase/SnRK1-Activating Kinase (GRIK/SnAK) proteins differentially regulate salt and energy stress responses in Arabidopsis

**DOI:** 10.1101/2023.03.10.532032

**Authors:** Nathalie Crepin, Filip Rolland

**Affiliations:** Laboratory of Molecular Plant Biology, Plant Metabolic Signaling, Biology Department, KU Leuven, Kasteelpark Arenberg 31, 3001 Heverlee-Leuven, Belgium; KU Leuven Plant Institute (LPI), Kasteelpark Arenberg 31, 3001 Heverlee-Leuven, Belgium

**Keywords:** GRIK/SnAK, SnRK1, metabolic stress, SOS2, salt stress, S-acylation, tonoplast localization

## Abstract

SnRK1 and SnRK3.11/SOS2 are key protein kinases in plant cellular energy and salt stress signaling, respectively. Using cellular assays, we confirm that the GRIK/SnAK (Geminivirus Rep-Interacting Kinase/SnRK1-Activating Kinase) proteins act as their main activating upstream kinases in Arabidopsis, catalyzing T-loop phosphorylation on the SnRK1α1 T175 and SOS2 T168 residues. Remarkably, SnRK1α1 phosphorylation on the neighbouring S176 residue competes with GRIK-mediated T175 phosphorylation to negatively regulate SnRK1 activity. Cellular assays and transgenic plants also revealed that the GRIK proteins, via N-terminal S-acylation, are predominantly localized at the tonoplast, where they interact with SnRK1α1 and SOS2. We optimized a leaf mesophyll protoplast-based Acyl PEG Exchange (APE) protocol to further explore GRIK protein S-acylation and tonoplast recruitment and identified the amino acid residues involved. GRIK1 tonoplast localization is likely mediated by initial membrane sampling via N-terminal domain hydrophobicity and local S-acylation, independently of a secretory pathway. Finally, *grik1-1 grik2-1* double KO mutants complemented with a non-S-acylatable mutant GRIK1 protein exhibit increased salt sensitivity (reduced SOS2 activity) but hyperactive SnRK1 signaling, demonstrating the differential importance of GRIK subcellular localization for Arabidopsis salt and energy stress responses.

## Introduction

The GRIK/SnAK (Geminivirus Rep-Interacting Kinase/SnRK1-Activating Kinase) proteins (GRIK1/SnAK2 and GRIK2/SnAK1, respectively) were originally identified in a yeast-2-hybrid screen of an Arabidopsis seedling cDNA library using the replication-associated geminiviral Rep/AL1 protein of the Tomato Golden Mosaic Virus (TGMV) as bait (Kong and Hanley-Bowdoin, 2002). Follow-up studies found that both GRIK proteins can functionally complement a yeast strain devoid of the three upstream activating kinases of Snf1 (Sucrose non-fermenting1), the fungal ortholog of plant SnRK1 (Snf1-Related Kinase1) and animal AMPK (AMP-activated kinase) (Shen and Hanley-Bowdoin, 2006; Hey et al., 2007). The highly conserved eukaryotic SNF1/SnRK1/AMPK kinases act as cellular fuel gauges, integrating and responding to diverse environmental and developmental cues that affect energy status. They typically inhibit energy consuming biosynthetic and growth processes, while activating catabolic reactions to maintain energy homeostasis, both through direct targeting of key metabolic enzymes and transcriptional regulation (Broeckx et al., 2016; Hulsmans et al., 2016; Crepin and Rolland, 2019). These kinases function as heterotrimeric complexes, with a catalytic α subunit (SnRK1α1/KIN10 and SnRK1α2/KIN11 in Arabidopsis) and regulatory β and γ subunits (Broeckx et al., 2016). The GRIK kinases were found to catalyse phosphorylation of the SnRK1 catalytic α subunit at a highly conserved Thr residue (T175 in Arabidopsis SnRK1α1) in the activating T-loop, both *in vitro* (Hey et al., 2007; Shen et al., 2009; Crozet et al., 2010) and *in vivo* (Glab et al., 2017). This phosphorylation stabilises the T-loop in an open and extended conformation to facilitate substrate binding and is an absolute prerequisite for SNF1/SnRK1/AMPK kinase activity (McCartney and Schmidt, 2001; Hawley et al., 1996; Baena-Gonzalez et al., 2007). In turn, SnRK1 phosphorylates and inactivates the GRIK proteins, presumably as a negative feedback mechanism to avoid persistent signaling (Crozet *et al*., 2010). The GRIK proteins, like many kinases involved in innate immune signaling in both animals and plants, (Dardick and Ronald, 2006; Dardick *et al*., 2012; Bender *et al*., 2021), are non-RD kinases, lacking the typical arginine (R) residue preceding a highly conserved aspartate (D) residue responsible for correct positioning of the phospho-acceptor site of the substrate (Kornev *et al*., 2006) (**Suppl. Figure 1**). They also do not require T-loop phosphorylation to become active (Kornev et al., 2006).

The GRIK proteins were found to accumulate in infected tissue (Shen and Hanley-Bowdoin, 2006). Together with their targeting by viral pathogenicity factors, this indicates that downstream SnRK1-mediated metabolic reprogramming is part of the plant antiviral response (Hulsmans et al., 2016). However, GRIK proteins also distinctively accumulate in young developing tissue containing dividing and endoreduplicating cells, such as the shoot apical meristem (SAM), leaf primordia, flower buds and siliques (Shen and Hanley-Bowdoin, 2006), where sensing of metabolic/carbon status and energy supply is crucial. The lack of GRIK proteins in uninfected mature tissues was proposed to result from proteasomal degradation triggered by cell differentiation and maturation (Shen and Hanley-Bowdoin, 2006). In addition, the GRIK 5’ untranslated regions (UTRs) contain upstream open reading frames, encoding conserved peptides (CPuORFs), possibly involved in stress-responsive and cell and tissue type-specific GRIK translation (Shen and Hanley-Bowdoin, 2006; Jorgensen and Dorantes-Acosta, 2012; Van der Horst et al., 2020; Dever et al., 2020; Causier et al., 2022).

While Arabidopsis *grik1* and *grik2* single knock out (KO) mutants show a wild type (WT) phenotype under control conditions, suggesting functional redundancy, *grik1 grik2* double KO is lethal in soil-grown plants (Glab et al., 2017). These results were not unexpected as KO of both Arabidopsis SnRK1 catalytic α subunits is also lethal, and virus-induced gene silencing (VIGS) of both α subunits results in a severely stunted growth and eventually death (Baena-Gonzalez et al., 2007). However, *grik1 grik2* lethality could be bypassed by growing seedlings *in vitro* on 3% glucose. These conditions enabled confirmation that *grik1 grik2* double mutants lack SnRK1 activity coinciding with stunted growth and sterility, producing round flower buds with protruding stigmata (Glab et al., 2017). It was concluded that *grik1 grik2* mutants fail to become autotrophic and require continuous exogenous sugar supply for survival (Glab et al., 2017).

In addition to the conserved heterotrimeric SnRK1 proteins, plants also evolved monomeric SnRK2 kinases, notably involved in ABA (abscisic acid)-mediated stress signaling, and SnRK3 or calcineurin B-like (CBL)-interacting protein kinases (CIPKs) (Hrabak et al., 2003). The SnRK (SnRK1 and plant-specific SnRK2 and SnRK3) kinases function in an intricate nutrient and stress responsive network with the antagonistic growth-promoting TOR (Target Of Rapamycin) kinase (Crepin and Rolland, 2016; Belda-Palazon et al., 2022). Interestingly, GRIK1 was found to also phosphorylate the salt stress signaling SnRK3.11/CIPK23/SOS2 (SALT OVERLY SENSITIVE2) kinase (Liu et al., 2000) *in vitro* at the activating T168 residue (Barajas-Lopez et al., 2018). To cope with a pronounced rise in cytosolic Na^+^ upon salt stress, plants use three mechanisms: (i) minimizing Na^+^ import into the cell, (ii) Na^+^ sequestration into the vacuole and, (iii) Na^+^ extrusion out of the cell (Ji et al., 2013). When Na^+^ stress is perceived, plant cells immediately respond by increasing cytosolic Ca^2+^ levels (Knight et al., 1997). In root cells specifically, the Ca^2+^ signal is recognized by the Ca^2+^-binding SOS3/CBL4 protein, that then interacts with the autoinhibitory domain of SOS2, resulting in the formation of a SOS2-SOS3 protein complex and activation of the SOS2 kinase (Halfter et al., 2000). This protein complex is subsequently recruited to the plasma membrane where SOS2 phosphorylates and activates the SOS1 Na^+^/H^+^-antiporter that exports Na^+^ from the cytosol into the extracellular space (Qiu et al., 2002; Ji et al., 2013). In shoot cells, however, another pathway is activated, involving SOS2 and CBL10. Similar to SOS3, CBL10 activates SOS2 by interaction, but the SOS2-CBL10 complex is predominantly targeted to the tonoplast, where it promotes Na^+^ import into the vacuole (Kim et al., 2007; Quan et al., 2007; Yang et al., 2019) by activating the vacuolar Na^+^/H^+^-exchanger (NHX) (Qiu et al., 2004) and/or vacuolar H^+^-ATPases (V-ATPases) (Batelli et al., 2007; Krebs et al., 2010). SOS2 was also reported to activate the tonoplast Ca^2+^-exchanger CAX1, providing a link between Na^+^-and Ca^2+^-homeostasis in plants (Cheng et al., 2003). However, none of the latter three targets were demonstrated to be directly phosphorylated by SOS2 (Qiu et al., 2004; Batelli et al., 2007).

SOS2 can thus be activated in two ways: through interaction with a CBL protein, relieving SOS2 auto-inhibition (Halfter et al., 2000), or via GRIK1-mediated T-loop phosphorylation at T168 (Guo et al., 2001; Barajas-Lopez et al., 2018). These mechanisms are not mutually exclusive and when combined result in SOS2 hyperactivity (Guo et al., 2001; Qiu et al., 2002). In the absence of SOS3, a constitutively active SOS2-T168D mutant could phosphorylate SOS1 *in vitro* (Guo et al., 2001). However, SOS2-T168D complementation of a *sos3* KO mutant could not rescue the root salt hypersensitivity (Guo et al., 2001). Hence, to elicit an effective salt stress response, not only kinase activity, but also correct SOS2 subcellular localization is important. The latter is determined by the lipid modification of the CBL that binds to the SOS2 protein. For example, plasma membrane localization of SOS2 is mediated by the N-terminally myristoylated SOS3 (Ishitani et al., 2000), whereas SOS2 tonoplast targeting is mediated by the S-acylated CBL10 (Chai et al., 2020).

N-myristoylation and S-acylation are two types of lipid modifications that direct proteins to intracellular membranes. N-myristoylation typically occurs co-translationally on a N-terminal Gly residue after removal of the initiating Met, requires a specific established consensus motif, and is irreversible (Resh, 2013). S-acylation generates a stronger membrane affinity and is unique due to its reversibility that allows for dynamic regulation (Chen et al., 2021). A recent screen revealed that 6% of the Arabidopsis proteome is S-acylated, suggesting an important role for this lipid modification in various cellular functions, such as signaling, trafficking, and metabolism (Kumar et al., 2022). Although protein S-acylation can also occur spontaneously (Guan and Fierke, 2011; Li and Qi, 2017), it is generally catalyzed by protein S-acyl transferases (PAT). These are integral membrane proteins containing a highly conserved Asp-His-His-Cys (DHHC) catalytic motif, in which the Cys (C) residue is essential for enzymatic activity (Zhou et al., 2013). PATs covalently attach a fatty acid (commonly palmitic acid) to a cysteine residue of the substrate protein via a labile thioester bond (Schmidt, 1989). The Arabidopsis genome encodes 24 DHHC-containing PATs localized in various intracellular membranes, including the plasma membrane and membranes of the Golgi apparatus, ER and vacuole, implying that S-acylation can happen at any stage of the secretory pathway or at the destination membrane (Batistic, 2012).

S-acylation can affect protein stability, promote protein-protein interactions or trigger conformational changes that modulate enzyme activity (Chen et al., 2021) but its major function is to increase protein hydrophobicity and hence membrane affinity (Hemsley and Grierson, 2008; Li et al., 2022). However, a recent report that the SOS3 protein requires S-acylation for plasma membrane release and subsequent nuclear translocation (Villalta et al., 2021) indicates that its effects are more complex. Removal of the acyl group or de-acylation by hydrolysis of the labile thioester linkage is mediated by thioesterases. Only recently, the ABAPT (Alpha/Beta hydrolase domain-containing protein 17-like acyl protein thioesterase) proteins were identified as putative de-S-acylating enzymes in Arabidopsis (Liu et al., 2021). While many proteins are more stably S-acylated to modulate association with the target membrane, others undergo rapid cycling of S-acylation/de-acylation as a switch mechanism between initiation and termination of signaling events (Li et al., 2022).

Here, we first confirm *in vivo* GRIK1-mediated activating phosphorylation of the SnRK1α1 T175 and SOS2 T168 T-loop residues and investigate additional regulatory SnRK1α1 phospho-sites. We further demonstrate that the GRIK proteins are predominantly localized at the tonoplast via N-terminal S-acylation and that they recruit SnRK1α1 and SOS2 to this membrane for T-loop phosphorylation. An optimized leaf mesophyll protoplast-based Acyl PEG Exchange (APE) protocol can (i) detect the number of S-acylation events and, when combined with mutational analysis, (ii) identify the S-acylated cysteine residues of the protein of interest. Finally, complementation of *grik1-1 grik2-1* double KO mutants with a non-S-acylatable GRIK1 protein reveals the differential importance of GRIK1 subcellular localization for Arabidopsis energy and salt stress responses.

## Results

### GRIK1 phosphorylates the SnRK1α1 and SOS2/SnRK3.11/CIPK24 activating T-loop residues *in vivo*

Eukaryotic RD kinases require phosphorylation of a specific amino acid residue (Ser/Thr/Tyr) located in their T-loop to render the protein catalytically competent (Taylor and Kornev, 2011). The GRIK proteins were identified as major upstream kinases of SnRK1α1/KIN10 and SOS2/SnRK3.11/CIPK24. More specifically, GRIK1 was shown to activate SnRK1α1 (Shen et al., 2009; Crozet et al., 2010; Zhai et al., 2018) and SOS2 (Barajas-Lopez et al., 2018) *in vitro* by phosphorylation of the T-loop residues T175 and T168, respectively. While GRIK1-mediated phosphorylation of SnRK1α1 T175 was also confirmed *in planta* (Glab et al., 2017), *in vivo* evidence for SOS2 T168 phosphorylation by GRIK1 is still lacking. Hence, we set out to further investigate both SnRK1α1 and SOS2 T-loop phosphorylation by GRIK1 using cellular assays.

First, we transiently overexpressed SnRK1α1 with or without GRIK1 in Arabidopsis leaf cells, including the kinase-dead SnRK1α1-K48M and GRIK1-K137A mutant proteins as negative controls (Baena-Gonzalez et al., 2007; Shen et al., 2009), to confirm SnRK1α1 T-loop phosphorylation using a Phos-tag acrylamide-based mobility shift assay (Kinoshita et al., 2009; Ramon et al., 2019). The previously reported significant SnRK1α1 T-loop (T175) phosphorylation without GRIK co-expression (Baena-Gonzalez et al., 2007; Shen et al., 2009; Ramon et al., 2019) is reduced in the kinase-dead SnRK1α1-K48M mutant protein (**Figure 1A**). This is indicative of SnRK1α1 T-loop autophosphorylation and consistent with the proposed default active state of SnRK1α1 (Ramon et al., 2019; Crepin and Rolland, 2019). The increased protein levels observed for SnRK1α1-K48M and SnRK1α1-T175A is consistent with SnRK1 feedback triggering its own degradation (Crozet et al., 2016). Co-expression of WT GRIK1, but not the kinase-dead GRIK1-K137A mutant protein, increased T175 phosphorylation of both SnRK1α1 and SnRK1α1-K48M, confirming that GRIK1 phosphorylates SnRK1α1 at T175 *in vivo*. We then investigated GRIK1-mediated SOS2 T168 phosphorylation using a similar approach. Without GRIK1 co-expression, SOS2 produced several protein bands in the Phos-tag immunoblot, indicative of phosphorylation (**Figure 1B**). Co-expression of GRIK1 resulted in two extra protein bands. These bands did not appear with the SOS2-T168A mutant protein (**Figure 1B**), indicating that they are associated with T168 phosphorylation and that the SOS2 T168 residue is phosphorylated by GRIK1 *in vivo*. These results also suggest that, unlike SnRK1α1, SOS2 is not T-loop phosphorylated under control conditions. Since GRIK1 phosphorylates both SnRK1 and SnRK3 proteins, we wondered whether it can also target SnRK2 proteins. However, SnRK2.3 did not produce additional phospho-bands in a Phos-tag experiment with GRIK1 co-expression (**Figure 1C**), consistent with a previous report that GRIK1 failed to phosphorylate SnRK2.4 (Shen and Hanley-Bowdoin, 2006). Possibly, RAF-like kinases act as the dominant or exclusive SnRK2 upstream kinases (Lin et al., 2021; Maszkowska et al., 2021).

**Figure 1:**
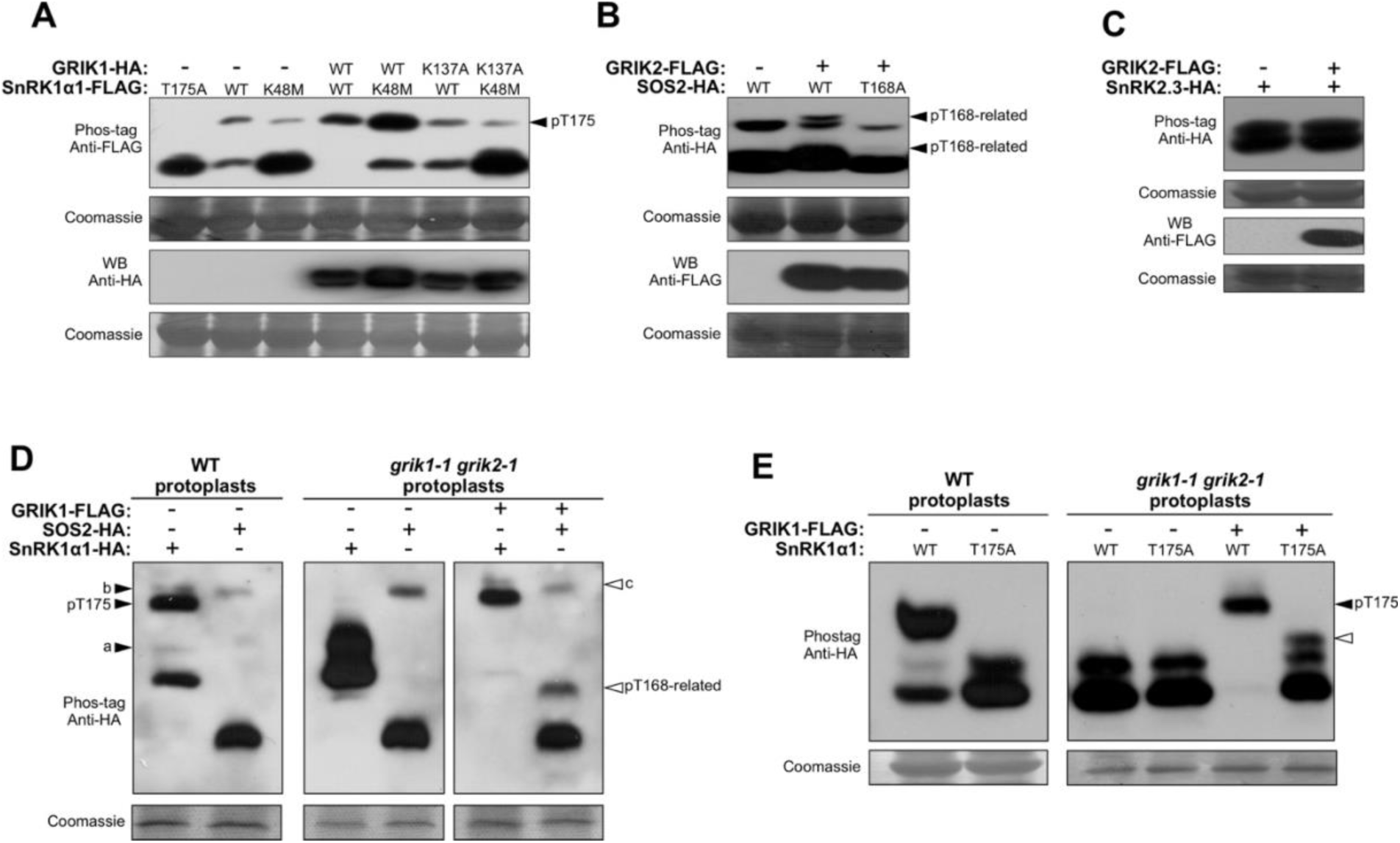
In vivo SnRK1α1 and SOS2 phosphorylation by GRIK1. Phos-tag acrylamide-based mobility shift assays using transient expression of HA-tagged WT or mutant SnRK1α1 (A), SOS2 (B) and SnRK2.3 (C) without and with FLAG-tagged GRIK1 in wild type (A-D) or grik1-1 grik2-1 mutant (D,E) leaf mesophyll protoplasts. SnRK1α1-K48M and GRIK1-K137A mutant proteins serve as kinase-dead negative controls. pT175 and pT168-related indicate phospho-bands associated with T175-phosphorylated SnRK1α1 and T168-phosphorylated SOS2, respectively. Letters “a” and “b” indicate bands resolved due to optimized Phos-tag experimental settings. The letter “c” indicates a GRIK1-independent SOS2 phospho-band. Phos-tag resolving gels contained 7.5% acrylamide, 25 μM Phos-tag and 50 μM MnCl_2_ (A,B,C) or 10 % acrylamide, 50 μM Phos-tag and 100 μM MnCl_2_ (D,E). Immunoblot analyses were performed using anti-HA and anti-FLAG antibodies. Coomassie blue staining of the large subunit of Rubisco served as loading control.

Remarkably, GRIK protein expression was reported to be restricted to young developing tissues or virus-infected mature tissue (Shen and Hanley-Bowdoin, 2006; Shen et al., 2009). This suggests the presence of alternative SnRK1α-activating kinases (Shen and Hanley-Bowdoin, 2006; Shen et al., 2009). We explored this further using a *grik1-1 grik2-1* double mutant. The inability of this mutant to become autotrophic can be bypassed by *in vitro* growth on glucose medium (Glab et al., 2017). We crossed the single *grik1-1* (GABI_713C09) and *grik2-1* (SALK_015230) T-DNA insertion lines (**Suppl. Figure 2A-B**) and isolated homozygous *grik1-1 grik2-1* double mutants from the progeny of *grik1-1 (+/-) grik2-1 (-/-)* sesquimutants that retained one WT *GRIK1* allele (Glab et al., 2017). Progeny of the alternative *grik1-1 (-/-) grik2-1 (+/-)* sesquimutants never produces the double mutant genotype. Growth for 4 weeks on half-strength (1/2) MS medium supplemented with 3% glucose in a 12 h light photoperiod readily distinguishes *grik1-1 (-/-) grik2-1 (-/-)* seedlings by their small size (**Suppl. Figure 2C**). After genotype confirmation by PCR, homozygous plantlets were transferred to plant culture boxes for *in vitro* growth for an additional two months under the same growth conditions to produce sufficient leaf material for mesophyll protoplast isolation. WT Arabidopsis plants were grown alongside (**Suppl. Figure 2C**).

We then transiently expressed SnRK1α1 and SOS2 without or with GRIK1 in WT and *grik1-1 grik2-1* mutant leaf cells to examine possible non-GRIK-mediated phosphorylation. The Phos-tag SDS-PAGE experimental settings were optimized for the SnRK1α1 and SOS2 proteins to enhance phospho-band separation and resolution. In these assays, SnRK1α1 was no longer phosphorylated at T175 when expressed in the *grik1-1 grik2-1* mutant background, while GRIK1 co-expression restored the phospho-band associated with pT175 (**Figure 1D**). This indicates that the GRIK proteins are the sole SnRK1α1-activating upstream kinases in mature Arabidopsis rosette leaves and that SnRK1α1 auto-phosphorylation depends on initial GRIK-mediated T175 phosphorylation, also implying that this occurs *in trans* rather than *in cis*. Optimized assays also enabled the discrimination of two additional weak SnRK1α1 phospho-bands in the WT background not observed previously (marked as “a” and “b” in Figure 1D). Intriguingly, phospho-band a, located just above the non-pT175 band, significantly increased in intensity in the *grik1-1 grik2-1* background, but completely disappeared with GRIK1 co-expression (**Figure 1D**). This indicates that the phosphorylation event is associated only with non-T-loop phosphorylated or inactive SnRK1α1. In contrast, phospho-band b immediately above the pT175 phospho-band is present in the WT background, absent in the *grik1-1 grik2-1* background and restored in the *grik1-1 grik2-1* background with GRIK1 co-expression (**Figure 1D**), suggesting a link with GRIK1-phosphorylated active SnRK1α1. In the optimized Phos-tag assay SnRK1α1-T175A co-expression with GRIK1 also produced an additional phospho-band, indicating that GRIK1 indeed phosphorylates other SnRK1α1 residues besides T175 (**Figure 1E**). The SOS2 phospho-band pattern, however, was identical in WT and *grik1-1 grik2-1* protoplasts (**Figure 1D**), suggesting that the upper SOS2 protein band (marked as “c”) is not associated with GRIK1 activity.

Combined, our data indicate that the GRIK proteins are the sole upstream kinases phosphorylating the activating T-loop residue of SnRK1α1 and possibly also SOS2.

### SnRK1α1 S176 phosphorylation competes with GRIK-mediated T175 phosphorylation

It was recently suggested that the GRIK proteins also phosphorylate the SnRK1α1 S176 residue adjacent to the activating T175 residue, thereby inhibiting SnRK1 activity (Martinez-Barajas and Coello, 2020). This would imply that the GRIK proteins act both as SnRK1α1-activating and SnRK1α1-inactivating kinases. We therefore examined whether and how SnRK1α1 S176 phosphorylation affects SnRK1α1 activity. We generated the SnRK1α1-S176A and phospho-mimetic SnRK1α1-S176D mutant proteins and examined their ability to induce *DIN6 (DARK INDUCED6)* promoter activity, a physiologically relevant readout of *in vivo* SnRK1 activity (Baena-Gonzalez et al., 2007; Dietrich et al., 2011). As expected, transient expression of WT SnRK1α1 enhanced activity of a luciferase (LUC) reporter under control of the *DIN6* promoter (*prDIN6::LUC)* (**Figure 2A**). However, expression of the SnRK1α1-S176D protein resulted in lower induction of *DIN6* promoter activity (**Figure 2A**). This suggests that S176 phosphorylation might indeed be a novel regulatory mechanism negatively affecting SnRK1α1 signaling. SnRK1α1-induced *prDIN6* activity requires nuclear SnRK1α1 localization (Ramon et al., 2019) and confocal fluorescence microscopy showed that SnRK1α1-S176D-GFP is still able to translocate to the nucleus (**Figure 2B**). However, Phos-tag analysis revealed a strongly reduced T175 phosphorylation (**Figure 2C**). Interestingly, GRIK1 co-expression partially restored SnRK1α1-S176D T175 phosphorylation (**Figure 2D**) as well as its ability to induce *DIN6* promoter activity (**Figure 2A)**. This is in contrast with the kinase-dead SnRK1α1-K48M protein, that also showed increased T175 phosphorylation with GRIK1 co-expression, but could not be re-activated (**Figure 2A,D**). Together, these data indicate that S176 phosphorylation negatively regulates SnRK1α1 activity by affecting phosphorylation of the activating T175 residue. This inhibition is partly relieved by GRIK1 overexpression, suggesting that other protein kinases compete with GRIK phosphorylation on adjacent residues to control SnRK1 activity.

**Figure 2:**
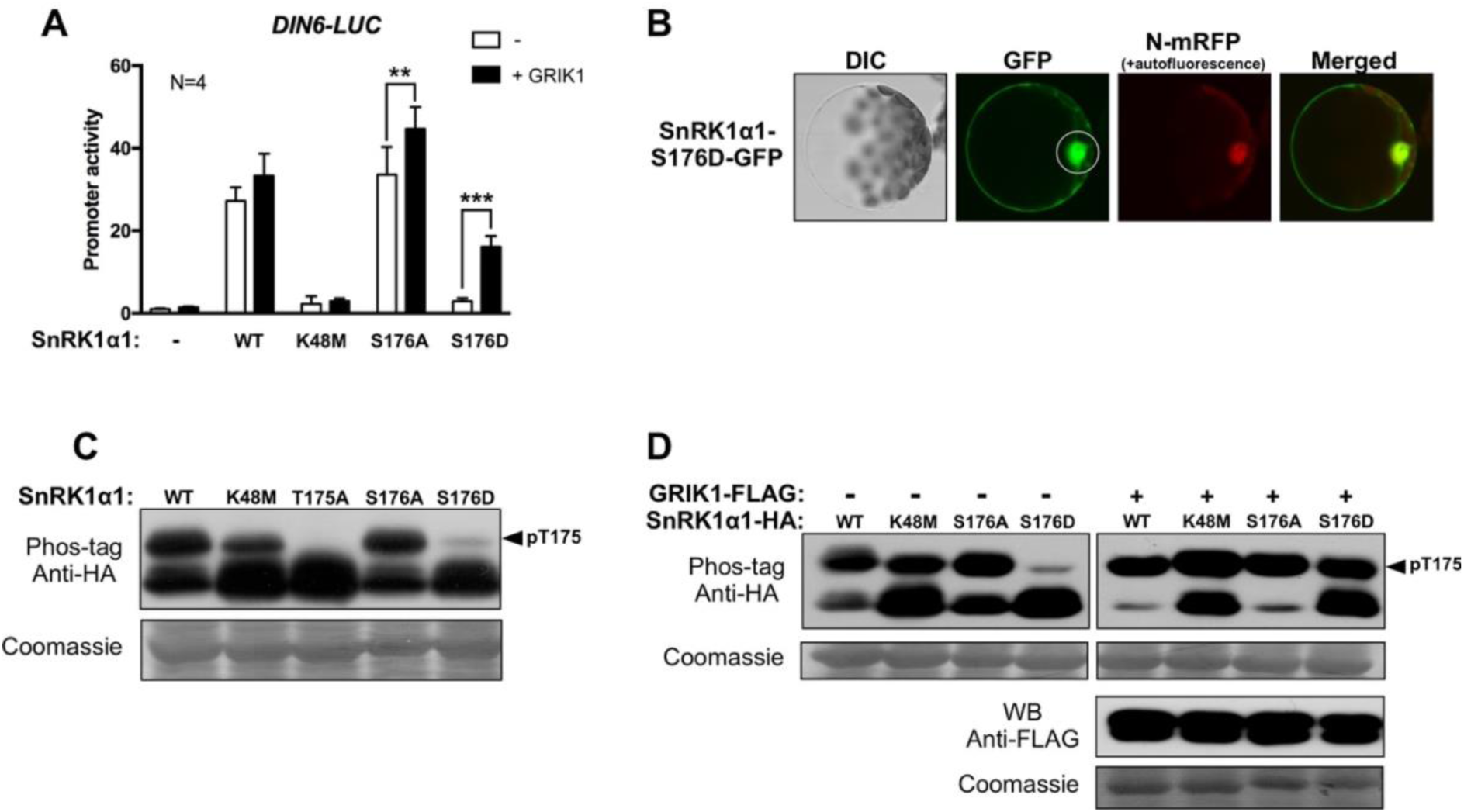
The SnRK1α1-S176D mutant protein shows compromised T175 phosphorylation, which can be partially rescued by GRIK1 co-expression. (A) DIN6 promoter activity in leaf mesophyll protoplasts transiently overexpressing WT or mutant (K48M, S176A or S176D) SnRK1α1 proteins without or with GRIK1. N=4 independent biological repeats (transfections). Statistical analysis was performed using Graphpad Prism 7, One-way ANOVA ***P<0.001, **P<0.01. (B) Confocal fluorescence microscopy of subcellular localization of the transiently expressed GFP-tagged SnRK1α1-S176D mutant protein. (C) Phos-tag mobility shift analysis of T175 phosphorylation of the WT and SnRK1α1-S176A and -S176D mutant proteins. SnRK1α1-K48M and SnRK1α1-T175A were added as negative controls for T175 phosphorylation and autophosphorylation activity, respectively. (D) Phos-tag mobility shift analysis of T175 phosphorylation of the WT SnRK1α1 and SnRK1α1-S176A and SnRK1α1-S176D mutant proteins transiently expressed in leaf mesophyll protoplast without or with GRIK1. SnRK1α1-K48M was added as a negative control for autophosphorylation activity. Immunoblot analyses of the Phos-tag mobility shift assays (resolving gel 7.5% acrylamide, 25 μM Phos-tag and 50 μM MnCl_2_) were performed using anti-HA and anti-FLAG antibodies. Coomassie blue staining of the large subunit of Rubisco serves as loading control.

### The GRIK proteins are localized and interact with SnRK1α1 and SOS2 at the tonoplast

We then explored the subcellular localization of the GRIK proteins and their interaction with the SnRK1α1 and SOS2 proteins using confocal fluorescence microscopy and bimolecular fluorescence complementation (BiFC) assays with GFP-and split YFP-tagged proteins in leaf mesophyll protoplasts (**Figure 3**). To our surprise, both GRIK proteins were localized predominantly at the vacuolar membrane (tonoplast) (**Figure 3A**). To further confirm tonoplast localization, a mild osmotic shock was applied in order to release vacuoles intactly (**Figure 3A**). Both GRIK-GFP fusion proteins also sporadically labelled small spherical structures located within the vacuole (**Figure 3B**). These tonoplast ‘bulbs’ are highly dynamic, multi-layered spherical membrane structures, generated by invagination and folding of the tonoplast, and are located in the vacuolar lumen (Saito et al., 2002). Moreover, we also detected GRIK1-GFP signals surrounding endocytosed chloroplasts (**Figure 3C**), corroborating tonoplast localization. While SnRK1α1 and SOS2 are nucleo-cytosolic proteins in control conditions (**Figure 3D**), BiFC assays indicate that their interaction with GRIK1 occurs at the tonoplast (**Figure 3E**). This suggests that GRIK1 recruits SnRK1α1 and SOS2 to the tonoplast for T-loop phosphorylation and activation.

**Figure 3.**
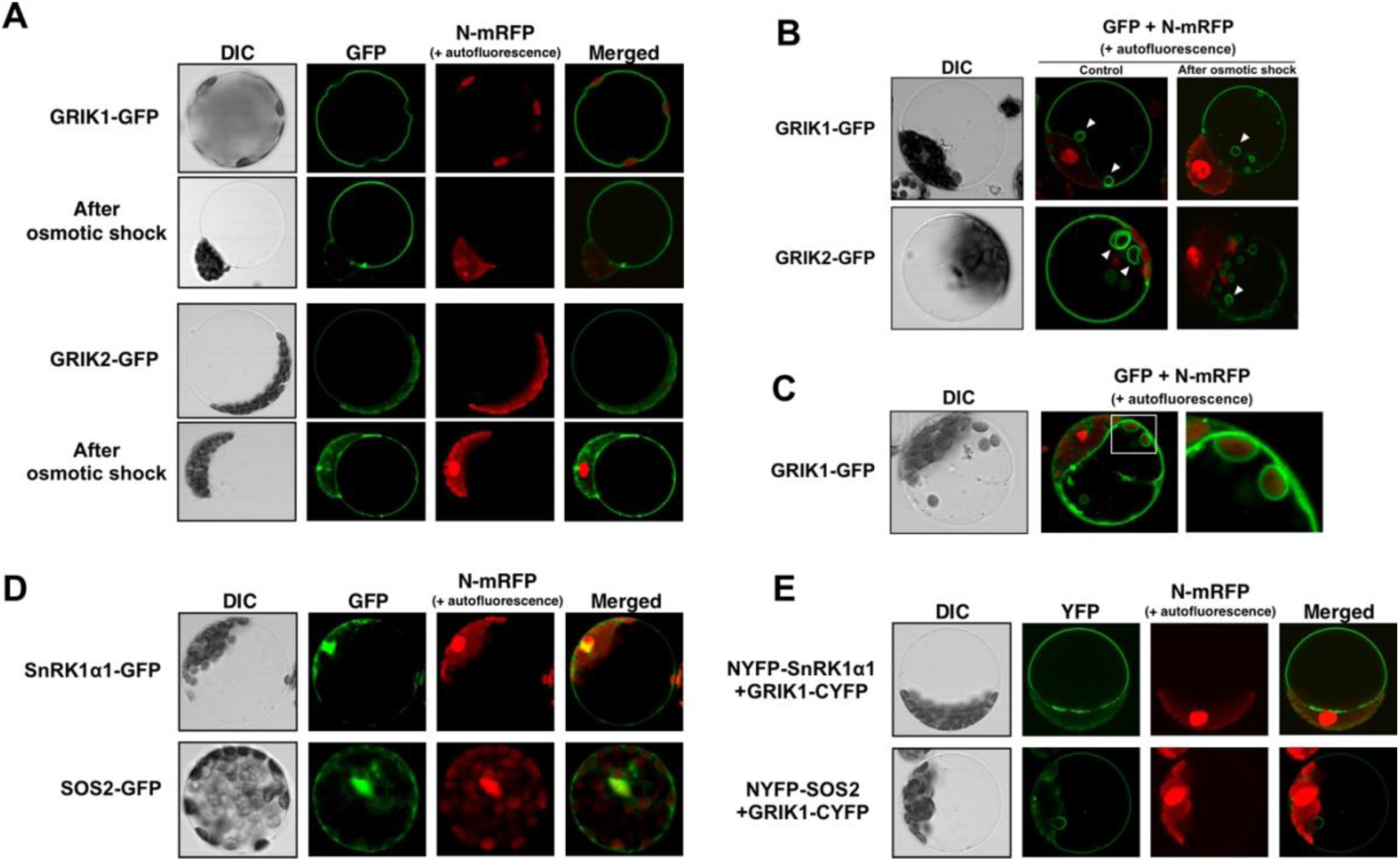
The GRIK proteins localise and interact with SnRK1α1 and SOS2 at the tonoplast. (A) Confocal fluorescence microscopy of subcellular localization of the transiently expressed GFP-tagged GRIK1 and GRIK2 proteins. (B) Tonoplast bulb labelling by GRIK1-GFP and GRIK2-GFP. Tonoplast bulbs are indicated with white arrowheads. (C) Endocytosed chloroplasts with GRIK1-GFP signals. The right image shows a magnification of the framed area in the middle image. (D) Confocal fluorescence microscopy of subcellular localization of transiently expressed GFP-tagged SnRK1α1 or SOS2. (E) Bimolecular fluorescence complementation (BiFC) analysis of the interaction between transiently expressed GRIK1 and SnRK1α1 or SOS2 in leaf mesophyll protoplasts. C-terminally tagged GRIK1-CYFP was co-expressed with N-terminally tagged NYFP-SnRK1α1 or NYFP-SOS2. Intact vacuoles were released using mild osmotic shock to highlight tonoplast localization. mRFP-tagged SCF30 (Splice Factor 30) was used as a nuclear reporter (N-mRFP). DIC, differential interference contrast image.

### N-terminal S-acylation targets the GRIK proteins to the tonoplast

*In silico* analysis of the GRIK amino acid sequences using the DeepTMHMM online server (Hallgren et al., 2022) did not predict any transmembrane domains, suggesting that post-translational lipid modification might be facilitating GRIK tonoplast targeting. The GRIK amino acid sequences were further inspected for the presence of residues that might be N-myristoylated, prenylated, or S-acylated using different *in silico* predictors such as GPS-Lipid (Xie et al., 2016) and CSS-PALM (Ren et al., 2008). Both GRIK proteins lack N-myristoylation or prenylation motifs, but S-acylation prediction produced high scores for the N-terminal GRIK1 Cys3, Cys14 and Cys17 and for the GRIK2 Cys14 and Cys17 residues (**Figure 4A**). Consistently, treatment of leaf mesophyll protoplasts with 2-bromopalmitate (2-BP), a general inhibitor of protein S-acylation *in vitro* (Fukata et al., 2004) as well as *in vivo* (Webb et al., 2000), caused relocalization of transiently expressed GFP-tagged GRIK1 and GRIK2 proteins from the tonoplast to the cytosol and nucleus (**Figure 4B**). Since the C14 and C17 residues are conserved in both GRIK proteins and the C3 residue is exclusive to GRIK1 (**Figure 4A**), we further focussed on GRIK1 to determine the individual and combined contribution of these *in silico* predicted S-acylated residues to tonoplast localization. We generated single, double and triple GRIK1 Cys mutant alleles for transient overexpression in leaf mesophyll protoplasts. The GFP-tagged single GRIK1-C3A and -C17A, and the double GRIK1-C3A/C17A mutant proteins still showed significant tonoplast localization, as illustrated by the labelling of tonoplast bulbs (**Suppl. Figure 3**). Tonoplast targeting of the GRIK1-C14A single and GRIK1-C3A/C14A double mutant proteins appeared less efficient but was not completely lost as intact vacuoles still showed a GFP signal (**Suppl. Figure 3**). However, the GFP-tagged GRIK1-C14A/C17A double and GRIK1-C3A/C14A/C17A triple mutant proteins were no longer associated with the tonoplast, instead showing a nucleo-cytosolic localization (**Suppl. Figure 3**). These observations indicate that C14 is the most and C3 (not conserved in GRIK2) the least important residue for tonoplast targeting of GRIK1. We confirmed tonoplast localization of WT GRIK1 and loss of tonoplast localization of the GRIK1-C3A/C14A/C17A triple mutant protein (from here on also referred to as GRIK1-3C) *in planta* by generating transgenic Arabidopsis lines overexpressing the GFP-tagged WT GRIK1 (35S::GRIK1-GFP) or GRIK1-3C triple mutant protein (35S::GRIK1-3C-GFP) (**Figure 4C**).

**Figure 4:**
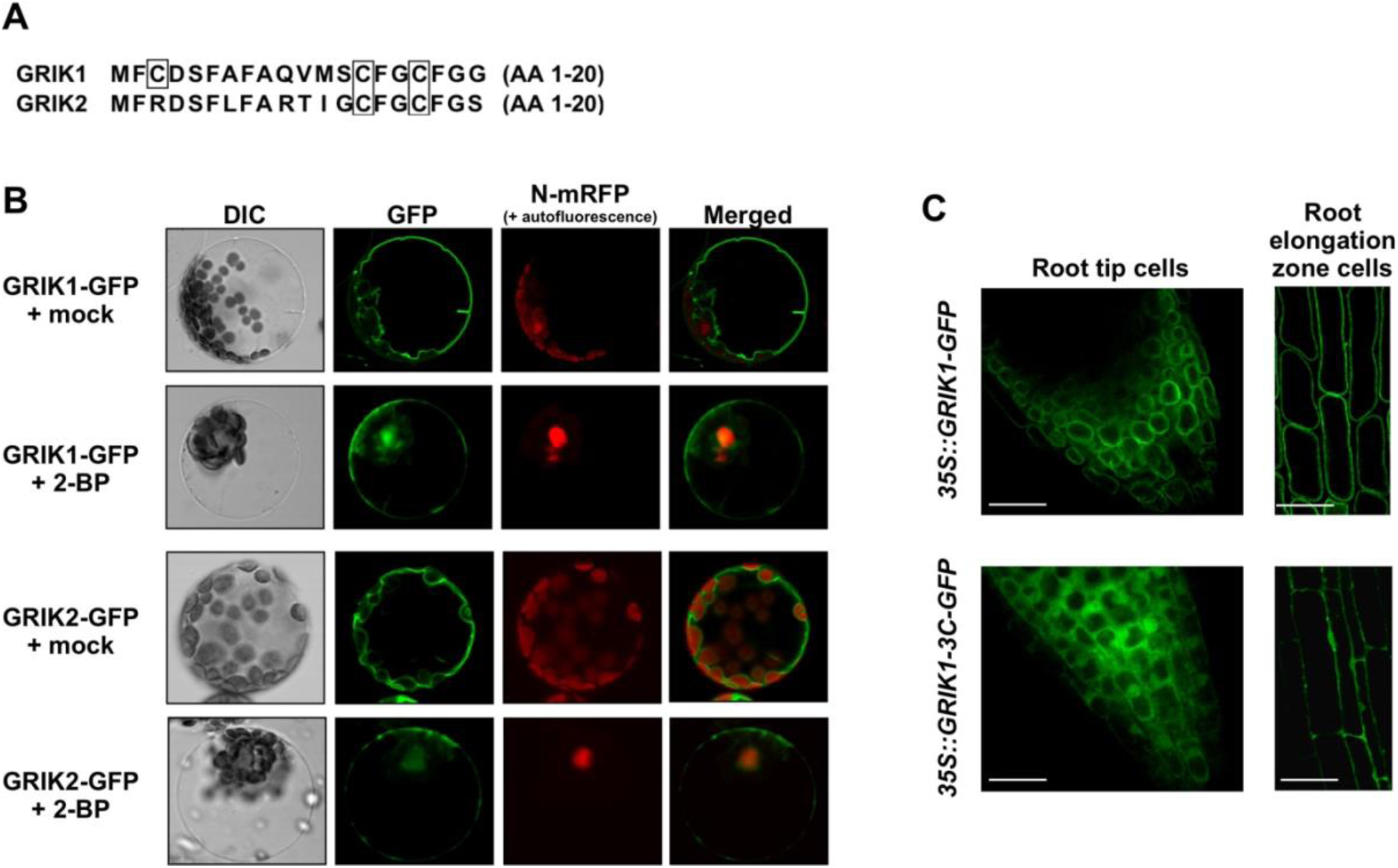
N-terminal S-acylation targets the GRIK proteins to the tonoplast. (A) Amino acid sequence alignment of the 20 N-terminal amino acid residues of the GRIK proteins. The in silico predicted putative S-acylated residues are framed. (B) Confocal fluorescence microscopy of subcellular localization of the transiently expressed GFP-tagged GRIK1 and GRIK2 proteins in leaf mesophyll protoplasts treated with either ethanol (mock) or 10 μM 2-BP. (C) Confocal fluorescence microscopy of transgenic Arabidopsis seedling roots overexpressing the GRIK1-GFP (35S::GRIK1-GFP) or GRIK1-3C-GFP (C3A/C14A/C17A) triple mutant protein (35S::GRIK1-3C-GFP). Scale bars indicate 25 μm.

We further investigated whether the GRIK1 N-terminal domain containing the S-acylated residues is sufficient for tonoplast targeting. We cloned the 20 N-terminal GRIK1 amino acid residues in frame with an HA-tagged enhanced green fluorescent protein (N20-eGFP/HA) and analyzed its subcellular localization. While WT eGFP/HA proteins were distributed throughout the nucleus and cytosol, the N20-eGFP/HA fusion protein was localized at the tonoplast and tonoplast bulbs (**Suppl. Figure 4**). Taken together, these results indicate that the GRIK1 20 amino acid N-terminal domain mediates tonoplast localization through S-acylation of the C3, C14 and C17 residues.

### An optimized protoplast-based Acyl PEG Exchange (APE) protocol confirms S-acylation of the Arabidopsis GRIK proteins

To confirm the results described above, we looked for an independent and complementary technique to identify S-acylation modifications of proteins. We started from an Acyl-PEG Exchange (APE) protocol used with mammalian cells for the identification of (the number of) S-acylated residues in proteins of interest through mass-tag labelling (Percher et al., 2016, 2017) and optimized the protocol for our experimental leaf mesophyll protoplast setup (**Figure 5A**). Cell lysates are first reduced with tris(2-carboxyethyl)phosphine (TCEP) to break any disulfide bridges and free thiol groups are blocked with N-ethylmaleimide (NEM). Next, thioester bonds between the acyl groups and cysteine residues are selectively cleaved with hydroxylamine (NH_2_OH) and the free thiol groups are alkylated with methoxy-PEG-maleimide (mPEG-Mal) of a specific mass. This results in a specific protein mass shift for each S-acylated cysteine residue, which can be visualized with SDS-PAGE and immunoblotting (**Figure 5A**). The number of mass shifts thus represents the number of S-acylated residues in the protein investigated. Specific modifications to the original APE procedure for Arabidopsis leaf mesophyll protoplast lysates can be found in Materials and Methods. To validate the optimized protoplast APE protocol, we used the Arabidopsis Calcineurin B-like protein CBL2. Similar to the GRIK proteins, CBL2 is S-acylated at the N-terminus, resulting in tonoplast localization (Batistic et al., 2012). CBL2 S-acylation is catalyzed by the tonoplast-localized PAT10 (Zhou et al., 2013), for which a confirmed KO mutant (*pat10-1*, SALK_024964) is available (Zhou et al., 2013). We first confirmed transiently expressed CBL2-GFP tonoplast localization and loss thereof in *pat10-1* mutant leaf mesophyll protoplasts (**Suppl. Figure 5A**). We then applied the APE protocol, using a 5 kDa mPEG-mal for the acyl-thiol exchange reaction, to determine the CBL2 S-acylation events in WT and *pat10-1* leaf mesophyll protoplasts using co-expression of CBL2 with WT PAT10 or an enzymatically inactive PAT10-C192A mutant protein (Zhou et al., 2013). The APE immunoblot produced two mass-shifted CBL2 protein bands in the WT background, confirming two previously reported S-acylated Cys residues (Batistic et al., 2012) (**Suppl. Figure 5B)**. Co-expression of PAT10 slightly increased the levels of S-acylated CBL2 (**Suppl. Figure 5B**), while PAT10-C192A co-expression had no effect at all, indicating that the catalytically inactive protein does not have the dominant-negative effect observed with mammalian protein S-acyl transferase (DHHC) proteins (Fukata et al., 2004; Fang et al., 2013; Lai and Linder, 2013). In the *pat10-1* background, only one very weak mass-shifted CBL2 protein band was detected without and with co-expression of the enzymatically dead PAT10-C192A mutant protein, while co-expression of the WT PAT10 restored CBL2 S-acylation to the levels observed in WT protoplasts (**Suppl. Figure 5B**). The remaining weak signal of the mono-S-acylated CBL2 variants is likely due to background labelling of non-cysteine nucleophiles with the mPEG-mal molecule (Percher et al., 2017). Alternatively, a PAT different from PAT10 also catalyzes CBL2 S-acylation, albeit insufficiently to target CBL2 to the tonoplast (**Suppl. Figure 5A**). Taken together, these data validate the modified APE protocol for determining the number of S-acylation events on proteins in Arabidopsis leaf mesophyll protoplasts.

**Figure 5:**
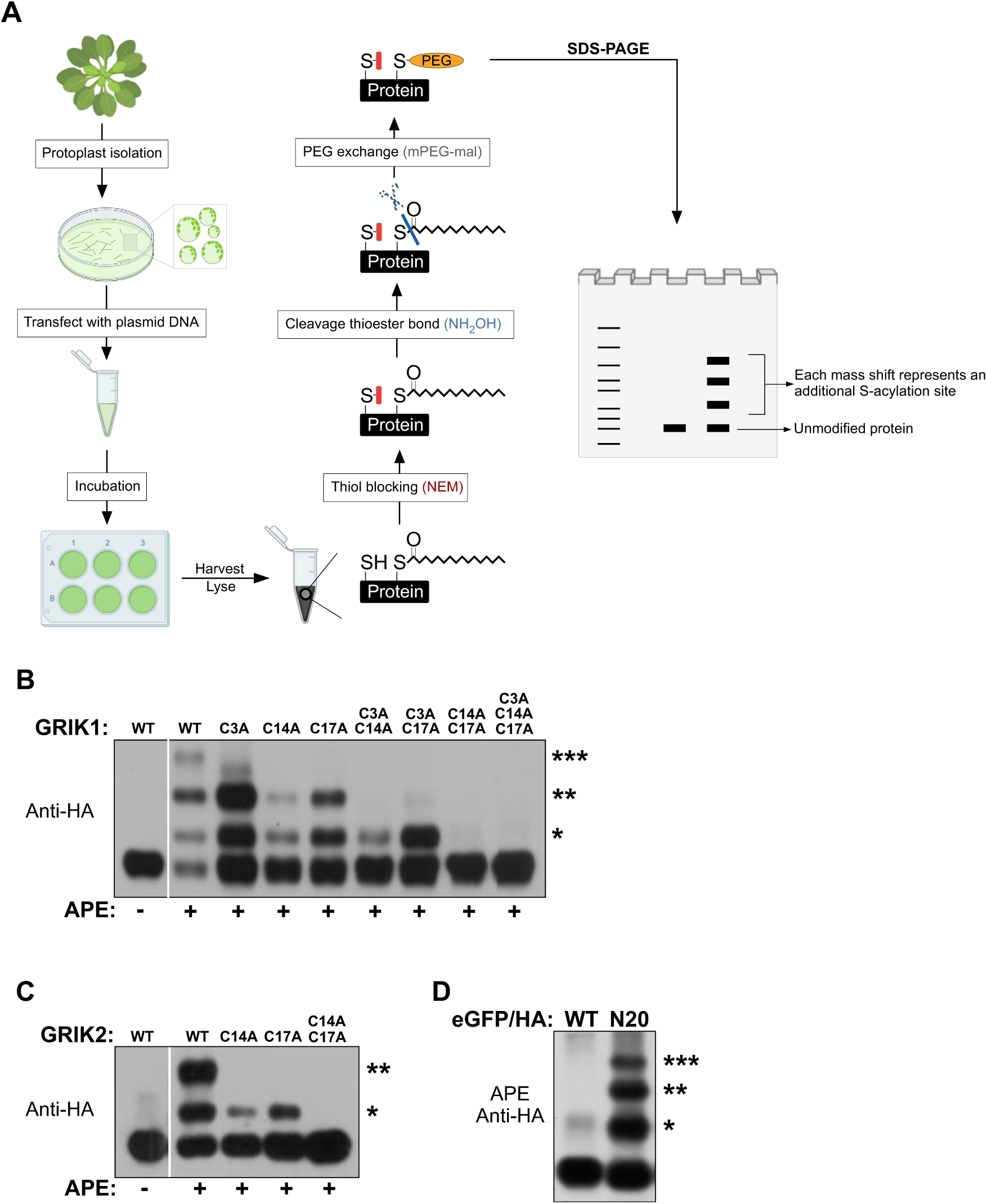
A protoplast-based Acyl PEG Exchange (APE) protocol confirms S-acylation of the Arabidopsis GRIK1 C3, C14 and C17 and of the GRIK2 C14 and C17 residues. (A) Workflow of the optimized assay. For detailed information, see text and “Materials and Methods”. Figure created with BioRender. (B-D) APE immunoblot assay of leaf mesophyll protoplasts transiently expressing HA-tagged WT, single, or combinatorial mutant GRIK1 proteins (B), WT, single, or double mutant GRIK2 proteins (C), and the WT eGFP/HA or N20-eGFP/HA fusion protein, containing the 20 N-terminal GRIK1 amino acids (D). Proteins were visualized using anti-HA antibodies. The number of PEGylations corresponds to the number of S-acylation events represented by asterisks.

We then applied our APE protocol to WT leaf mesophyll protoplasts transiently expressing GRIK1 or GRIK2. This confirmed respectively three and two S-acylation events for the respective WT proteins (**Figure 5B,C**). To assess whether the *in silico* predicted GRIK1 C3, C14 and C17 and GRIK2 C14 and C17 residues are the S-acylated residues, we repeated the APE experiment with the different single and combinatorial GRIK mutant proteins. With each substitution of a putative S-acylated Cys residue to Ala in the GRIK2 amino acid sequence, a mass-shifted GRIK2 protein variant disappeared from the APE immunoblot (**Figure 5C**). This confirms S-acylation of the GRIK2 C14 and C17 residues. Similarly, the GRIK1 single (C3A, C14A, C17A) and the double mutant (C3A/C14A and C3A/C17A) proteins lacked one and two S-acylation events respectively, while the GRIK1-3C triple mutant protein no longer showed any S-acylation, consistent with the GRIK1 C3, C14 and C17 residues being modified (**Figure 5B**). However, similar to the GRIK1-3C mutant protein, the double C14A/C17A mutant protein displayed no mass-shifts, even though the confirmed S-acylated C3 residue was still present. This is in agreement with our confocal microscopy results showing that the GRIK1-C14A/C17A mutant protein is no longer targeted to the tonoplast (**Suppl. Figure 3**). To further corroborate these results, we determined the number of S-acylation events on the N20-eGFP/HA fusion protein, only containing the 20 N-terminal GRIK1 residues. As expected, three shifted proteins band were observed (**Figure 5D**). Collectively, the optimized APE protocol successfully demonstrated S-acylation of the GRIK1 C3, C14 and C17 and the GRIK2 C14 and C17 residues in Arabidopsis leaf cells.

More careful inspection of the APE immunoblots revealed that GRIK1 C14A and GRIK2 C14A mutation decreases the efficiency of S-acylation of the remaining Cys residues (**Figure 5A,B**). In contrast, mutation of the GRIK1 C3 residue rendered S-acylation seemingly more efficient (**Figure 5A**). S-acylation has been shown to directly affect protein stability, increasing protein half-life (Chen et al., 2021). However, protein stability assays of the GRIK1-C3A and -C14A single mutant proteins, blocking *de novo* protein synthesis using the inhibitor cycloheximide (CHX) 4h after leaf mesophyll protoplast transfection, did not reveal any effect on GRIK1 protein stability (**Suppl. Figure 6**). Hence, these observations suggest a certain cooperativity between the different S-acylation sites.

### GRIK1 tonoplast localization is mediated by N-terminal domain hydrophobicity and independent of the secretory pathway

Because PATs are membrane proteins, initial transient membrane association of substrate proteins is essential to undergo S-acylation and subsequent stable membrane attachment (Hemsley and Grierson, 2008). N-myristoylation, prenylation, and transmembrane domains have all been reported as mechanisms for initial membrane binding of S-acylated proteins (Greaves and Chamberlain, 2006), but none of these motifs are present in the GRIK proteins. Transient membrane binding can also be accommodated via hydrophobic domains (Greaves and Chamberlain, 2006; Greaves et al., 2009; Konrad et al., 2014; Locatelli et al., 2020). Since efficient tonoplast targeting is established by the GRIK1 20 N-terminal amino acid residues, the signal that triggers initial transient membrane interaction must be present in that sequence. This peptide indeed consists of 50% hydrophobic in addition to 45% neutral and 5% acidic amino acids, rendering it highly hydrophobic (**Figure 6A**). Moreover, it includes five Phe (F) residues that are also conserved in GRIK2 (**Figure 6A**). If initial interaction of the GRIK1 protein with membranes is facilitated by its N-terminal domain, removal of its hydrophobicity should result in loss of membrane association and subsequent S-acylation. To verify this hypothesis, we generated GRIK1-F6A/F8A and GRIK1-F15A/F18A double mutant proteins and examined S-acylation. While mutation of the F6 and F8 residues had no effect, S-acylation of the GRIK1-F15A/F18A mutant protein was significantly affected (**Figure 6B**). This indicates that hydrophobicity is dispensable close to the GRIK C3 residue but crucial in direct proximity of the C14 and C17 residues for S-acylation to occur.

**Figure 6.**
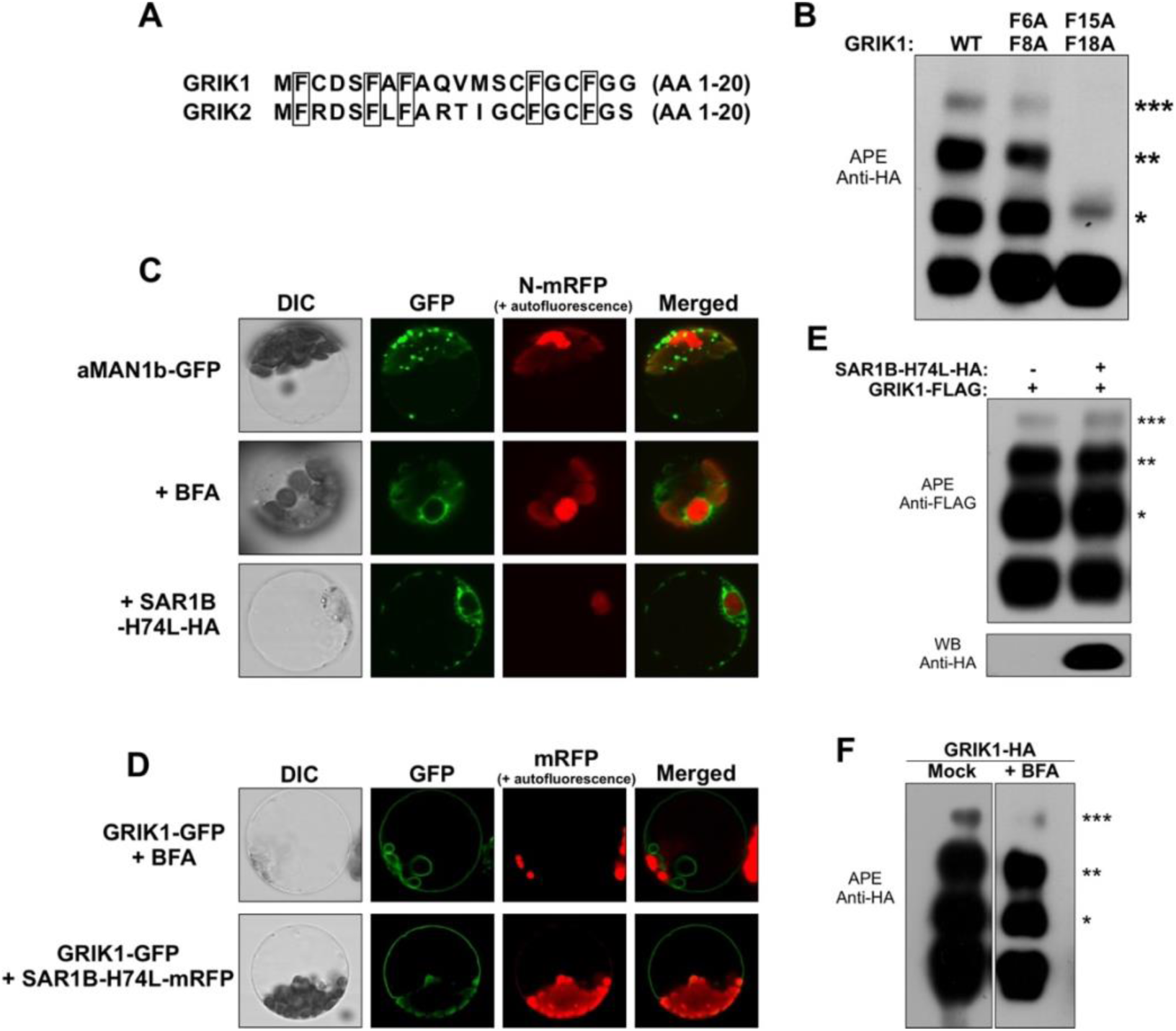
GRIK1 tonoplast localization is mediated by N-terminal domain hydrophobicity and independent of the secretory pathway. (A) Amino acid sequence alignment of the 20 N-terminal amino acid residues of the GRIK proteins. The highly conserved Phe (F) residues are framed. (B) APE immunoblot assay of leaf mesophyll protoplasts transiently expressing WT GRIK1 or mutant GRIK1-F6A/F8A or GRIK1 F15A/F18A mutant proteins. The number of PEGylations correspond to the number of S-acylation events represented by asterisks. Proteins were visualized using anti-HA antibodies. (C) Confocal fluorescence microscopy of the subcellular localization of transiently expressed GFP-tagged aMAN1b in leaf mesophyll protoplasts without or with BFA treatment of SAR1B-H74L-HA co-expression. (D) Confocal fluorescence microscopy of the subcellular localization of transiently expressed GFP-tagged GRIK1 with BFA treatment or SAR1B-H74L-mRFP co-expression. (E-F) APE immunoblot assay of leaf mesophyll protoplasts expressing FLAG-tagged WT GRIK1 without or with SAR1B-H74L co-expression (E) or BFA treatment (F). The number of PEGylations corresponds to the number of S-acylation events represented by asterisks. Proteins were visualized using anti-HA and anti-FLAG antibodies. BFA was added in the incubation medium to a final concentration of 50 μM immediately after transfection. mRFP-tagged SCF30 (Splice Factor 30) was used as a nuclear reporter (N-mRFP).

We hypothesize that the GRIK proteins use the weak membrane affinity of their N-terminal hydrophobic domain to sample various intracellular membranes. Upon contact with the membrane where their upstream PAT(s) reside(s), the GRIK proteins are S-acylated and subsequently stably attached to this membrane. The eventual delivery of the GRIK proteins to the tonoplast can be facilitated via two trafficking routes: one that uses components of the secretory pathway or one that directly targets to the tonoplast (Pedrazzini et al., 2013). A first indication that GRIK1 uses a direct route to the tonoplast, rather than the secretory pathway, comes from the observation that treatment with the S-acylation inhibitor 2-BP did not relocate GRIK1 to the endoplasmatic reticulum (ER), Golgi, or trans-Golgi network (TGN) (Batistic et al., 2012) (**Figure 4B**). To confirm this, we used two inhibitors that target different (COPI-and COPII-mediated) steps in the secretory pathway. The fungal toxin brefeldin A (BFA) inhibits formation of COPI vesicles, thereby promoting uncontrolled fusion of Golgi and ER membranes and trapping of proteins destined to other subcellular compartments (Nebenfurh et al., 2002). H74L mutation of the secretion-associated Ras1 (SAR1) GTPase protein that mediates ER-to-Golgi transport in Arabidopsis can be used to inhibit formation of COPII vesicles (Takeuchi et al., 2000). To confirm efficient BFA treatment and the dominant-negative effect of SAR1B-H74L mutant protein co-expression, we used the Golgi protein α-mannosidase 1b (αMAN1b) as a control (Kajiura et al., 2010). While BFA treatment and SAR1B-H74L co-expression resulted in relocalization of the αMAN1b protein from the Golgi apparatus to the ER (**Figure 6C**), GRIK1 was still observed at the tonoplast (**Figure 6D**), showing three S-acylation events (**Figure 6E,F**). This indicates that GRIK1 tonoplast targeting depends on a pathway which does not require either COPI or COPII vesicles. Based on these findings and the fact that 2-BP treatment results in GRIK1 nucleo-cytosolic localization, we conclude that GRIK1 moves from the cytosol to the tonoplast without use of secretory pathway vesicles.

### The PATs involved in GRIK S-acylation remain to be identified

A logic consequence of the model that the GRIK proteins first transiently interact with the tonoplast to which they are then stably attached after S-acylation, is that the PATs involved are tonoplast-localized. Two of the 24 putative Arabidopsis PATs, PAT10 and PAT11, are tonoplast proteins (Batistic, 2012). An APE immunoblot assay of *pat10-1* leaf mesophyll protoplasts transiently expressing the GRIK proteins still showed three and two S-acylation events for GRIK1 and GRIK2 respectively (**Suppl. Figure 7A**) and GRIK1 still showed significant tonoplast localization in *pat10-1* protoplasts (**Suppl. Figure 7B**). This indicates that PAT10 does not catalyze S-acylation of the GRIK proteins or that there is functional redundancy with other PAT proteins. Next, we isolated two homozygous *pat11* T-DNA insertion lines, *pat11-1* (SALK_022544) and *pat11-2* (SALK_026092) with T-DNA insertions at the start of the last exon (**Suppl. Figure 7C,D**). However, gene expression analysis indicates that neither is a complete KO at the transcriptional level (**Suppl. Figure 7E**). GRIK1 was also still targeted to the tonoplast in *pat11-1* mesophyll protoplasts (**Suppl. Figure 7F**).

### GRIK1 tonoplast targeting is important for salt stress responses

GRIK1-mediated SOS2 T-loop phosphorylation is essential for SOS2-mediated salt stress responses (Barajas-Lopez et al., 2018). To investigate the function of GRIK1 tonoplast localization *in planta*, we complemented *grik1-1 grik2-1* double KO by overexpressing either WT GRIK1 (*grik1-1 grik2-1* + WT) or the non-S-acylatable GRIK1-3C mutant protein (*grik1-1 grik2-1* + 3C). While no difference could be observed under control or osmotic stress (200 mM sorbitol) conditions, two independent mutant lines overexpressing GRIK1-3C (OE 26 and OE 35) germinated poorly on growth medium supplemented with 100 mM NaCl and germination was completely inhibited by 200 mM NaCl (**Figure 7A, Suppl. Figure 8A**). The mutant line complemented with WT GRIK1 also showed compromised germination compared to WT under these conditions, possibly because of an additional role for GRIK2 in the salt stress response. In addition, the effects of salt stress on cotyledon greening and root growth were examined by transferring 3-day-old seedlings from control conditions to 150 and 100 mM NaCl conditions respectively. GRIK1-3C complemented seedlings showed significantly more cotyledon bleaching (**Figure 7B, Suppl. Figure 8B**) as well as inhibition of root elongation (**Figure 7C, Suppl. Figure 8C**) compared to WT GRIK1 seedlings. Finally, we also measured expression of the salt-responsive genes *RD22* and *RD29a* (*RESPONSIVE TO DESSICATION 22* and *29a*) (Zhu, 2002) and *P5CS1* (encoding Delta1-pyrroline-5-carboxylate synthase 1, a key enzyme in proline biosynthesis) (Feng et al., 2016) using qRT-PCR. Consistent with the salt hypersensitive phenotype, induction of *RD22*, *RD29a* and *P5CS1* expression in seedlings in response to salt stress (300 mM NaCl for 4 h) was significantly lower in the *grik1-1 grik2-1* mutant complemented with GRIK1-3C (OE 35) than in WT seedlings and *grik1-1 grik2-1* mutants complemented with WT GRIK1 (**Figure 7D**). Combined, these data indicate that loss of GRIK1 tonoplast localization affects salt stress responses and increases salt sensitivity in Arabidopsis. Expression of the salt-response genes *RD29a* and *P5CS1* is controlled by the ETHYLENE AND SALT-INDUCIBLE ERF1 (ESE1) TF, in turn a target of the ETHYLENE INSENSITIVE3 (EIN3) TF (Zhang et al., 2011). Since SOS2 phosphorylates and activates EIN3 during salt stress (Quan et al., 2017), the reduced *RD29a* and *P5CS1* transcript levels in the *grik1-1 grik2-1* mutant complemented with GRIK1-3C is likely caused by compromised SOS2 kinase activity due to loss of T-loop (T168) phosphorylation. While mutant seedlings overexpressing WT GRIK1 showed elevated *ESE1* transcript levels compared to WT, both in control and 300 mM NaCl salt stress conditions, mutant seedlings overexpressing GRIK1-3C did not (**Figure 7E**). Consistently, in leaf mesophyll protoplasts transiently expressing SOS2, WT GRIK1 was more efficient at phosphorylating the SOS2 T168 residue than GRIK1-3C (**Figure 7F**). These data suggest that loss of GRIK1 tonoplast localization results in less efficient SOS2 T-loop phosphorylation and activation, required for appropriate target gene expression and growth and developmental responses to cope with salt stress.

**Figure 7:**
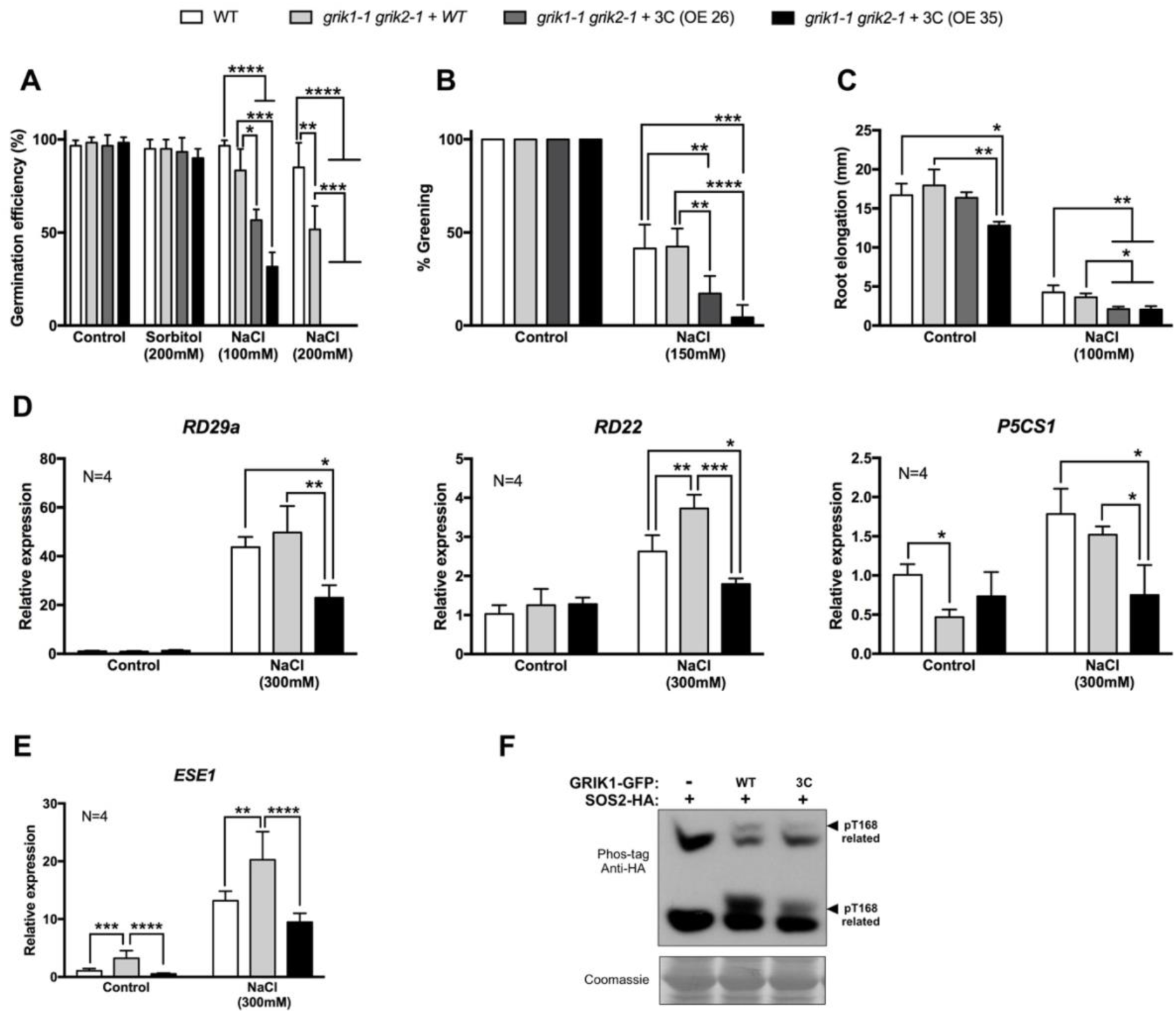
Loss of GRIK1 S-acylation and tonoplast localization affects Arabidopsis salt stress responses. WT and grik1-1 grik2-1 mutant lines complemented with WT GRIK1 or non-acylatable mutant GRIK1-3C were analysed for: (A) Germination efficiency on 1/2 MS medium supplemented with 1% sucrose without (control) or with 100 or 200 mM NaCl or 200 mM sorbitol (osmotic control) after 10 days in 16h light/8h dark conditions. Successful germination was defined as visible radicle protrusion. (B) Cotyledon greening 4 days after transfer of 3-day-old seedlings to fresh 1/2 MS growth medium supplemented with 1% sucrose without (control) or with 150 mM NaCl in 16h light/8h dark conditions. Green cotyledons were defined as not fully bleached. (C) Root elongation 3 days after transfer of 3-day-old seedlings to fresh 1/2 MS growth medium supplemented with 1% sucrose without (control) or with 100 mM NaCl in 16h light/8h dark conditions and turning the plate upside down for root bending. Root elongation was measured using the ImageJ software. N=3 with 20 seedlings per biological replicate. Data represent averages with standard deviations. Statistical analyses were performed with Graphpad Prism 7. One-way ANOVA, ****P<0.0001, ***P<0.001, **P<0.01, *P<0.1. Scale bars represent 0.5 cm. (D) RD29a, RD22 and P5CS1 expression in 7-day-old seedlings after transfer to liquid growth medium supplemented with 1% sucrose without (control) or with 300 mM NaCl for 4h. qRT-PCR analysis. Values are averages with standard deviations. N=4 with 10-15 pooled seedlings per biological replicate. Statistical analysis was performed using Graphpad Prism 7, One-way ANOVA, ***P<0.001, **P<0.01, *P<0.1. (E) ESE1 expression in 7-day-old seedlings after transfer to liquid growth medium supplemented with 1% sucrose without (control) or with 300 mM NaCl for 4 h. qRT-PCR analysis. Values are averages with standard deviations. N=4 biological repeats with 10-15 pooled seedlings per biological replicate. Statistical analysis was performed using Graphpad Prism 7, One-way ANOVA , **** P<0.0001, * P<0.1. Wild type leaf mesophyll protoplasts were used for (F) a Phos-tag mobility shift assay of transiently expressed SOS2-HA T168 T-loop phosphorylation without or with co-expression of WT GRIK1 or non-acylatable mutant GRIK1-3C. Bands associated with T168 phosphorylation are indicated with black arrowheads. Immunoblot analysis was performed using anti-HA antibodies. Coomassie blue staining of the large subunit of Rubisco served as loading control.

### Loss of GRIK1 tonoplast localization may enhance SnRK1 activation and target gene induction

Since GRIK-mediated T-loop phosphorylation is also required for SnRK1α1 activation, even under optimal conditions, we examined the importance of GRIK tonoplast localization for SnRK1 activity and signaling. In leaf mesophyll protoplasts, co-expression of the GRIK1-3C mutant protein appeared to be even more effective than co-expression of WT GRIK1 in enhancing SnRK1α1-induced *DIN6* target promoter activity (**Figure 8A**). A Phos-tag immunoblot assay confirmed that the GRIK1-3C mutant protein still efficiently phosphorylates the SnRK1α1 T175 residue (**Figure 8B**). We then assessed the effect on *in planta* SnRK1 signaling using a seedling sugar starvation assay and induction of *DIN6*, *BCAT2 (BRANCHED CHAIN AMINO ACID TRANSAMINASE2)* and *ATG8e (AUTOPHAGY8e)* target gene expression as readout (Ramon et al., 2019). *grik1-1 grik2-1* mutant seedlings complemented with WT GRIK1 induced a robust SnRK1 response, likely due to GRIK1 overexpression. Induction of *DIN6* and *ATG8e* was at least as strong in the mutant lines complemented with non-acylatable mutant GRIK1-3C 6h after sugar removal, and induction of *BCAT2* was even significantly stronger (**Figure 8C**). This is consistent with the protoplast assay results and suggests that loss of GRIK1 tonoplast localization may cause SnRK1α1 hyperactivation due to T-loop phosphorylation at different locations in the cell, no longer restricted to the tonoplast.

**Figure 8:**
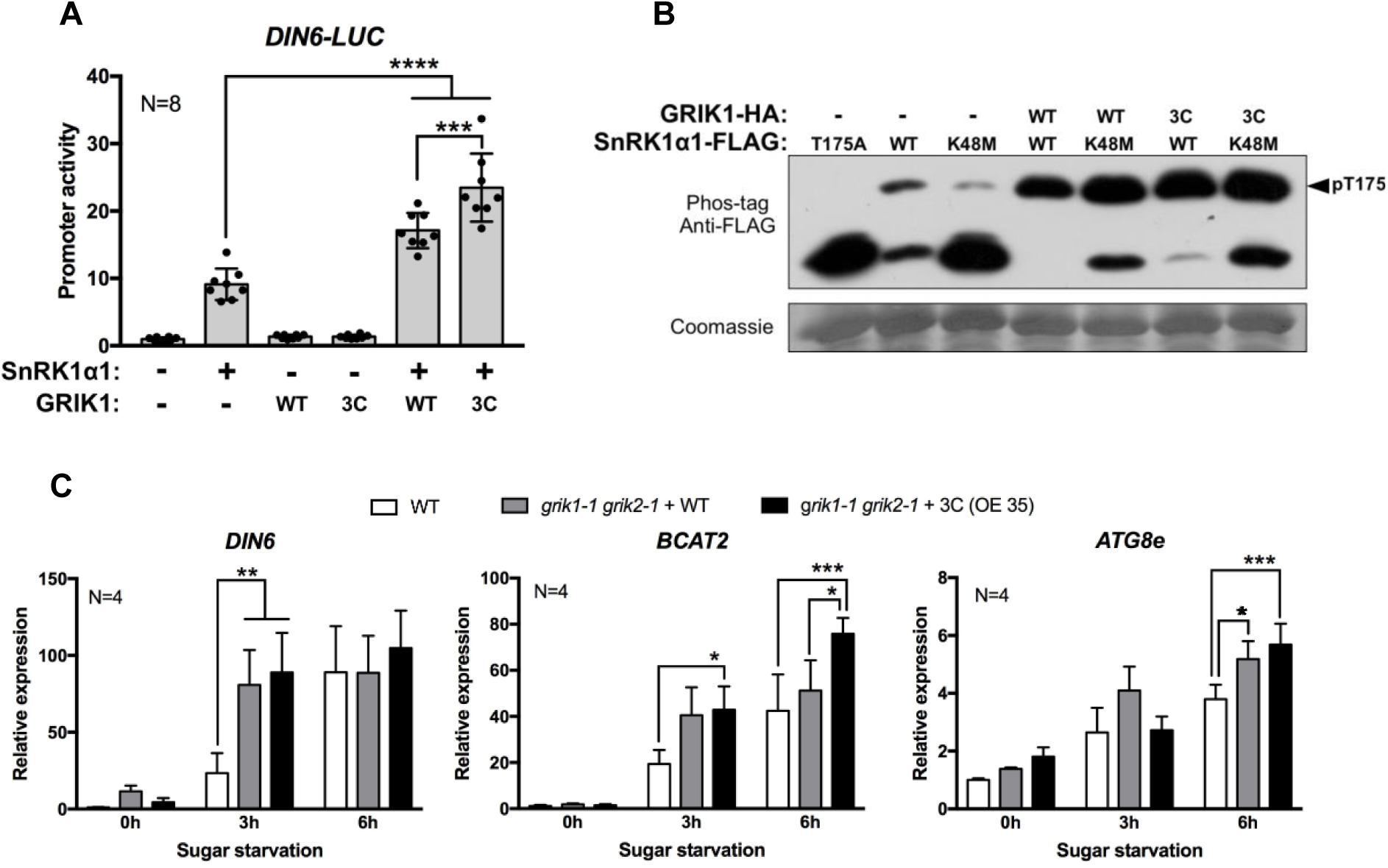
Loss of GRIK1 tonoplast localization enhances SnRK1 target gene activation. (A) DIN6 promoter activity in WT leaf mesophyll protoplasts upon transient expression of SnRK1a1 without or with WT GRIK1 or mutant GRIK1-3C. Results are averages of relative and normalized LUC values with standard deviations, N=8 independent biological repeats. Statistical analysis was performed using Graphpad Prism 7, One-way ANOVA ****P<0.0001, ***P<0.001. (B) Phos-tag immunoblot assay of transiently expressed SnRK1α1 T175 phosphorylation in leaf mesophyll protoplasts without or with co-expression of WT GRIK1 or mutant GRIK1-3C. The kinase-dead SnRK1α1-T175A and SnRK1α1-K48M mutant proteins served as a negative control for T175 phosphorylation and SnRK1α1 autophosphorylation, respectively. Proteins were visualized using anti-FLAG antibodies. Coomassie blue staining of the large subunit of Rubisco served as loading control. (C) qRT-PCR analysis of DIN6, BCAT2 and ATG8e expression in WT or grik1-1 grik2-1 double mutant seedlings complemented with either the WT GRIK1 protein or the GRIK1-3C mutant protein at different time points after transfer to liquid 1/2 MS devoid of sugars. Values are averages with standard deviations. N=4 biological repeats (10-15 seedlings pooled per repeat). Statistical analysis was performed using Graphpad Prism 7, One-way ANOVA, ***P<0.001, **P<0.01, *P<0.1.

## Discussion

The aims of this study were to further characterize the established but understudied GRIK protein kinases and to elucidate the significance of their role in regulating SnRK1 and SnRK3.11/SOS2 by T-loop phosphorylation and activation. We confirmed GRIK1-mediated phosphorylation of the activating T-loop residues T175 in SnRK1α1 and T168 in SOS2 *in vivo* using cellular and Phos-tag mobility shift assays. Our analyses show that a large fraction of the cellular SnRK1α1 protein pool is T-loop phosphorylated on T175, with significant autophosphorylation, and hence catalytically competent also under non-stress conditions. This is consistent with our hypothesis of a default SnRK1 activation and metabolic stress response with active repression via diverse mechanisms during optimal growth conditions with ample carbon and energy supplies (Ramon et al., 2019; Crepin and Rolland, 2019). In contrast, the SOS2 protein does not appear to be T168 phosphorylated under non-stress conditions, pointing to a significant difference in regulation by the upstream GRIK proteins.

Our analyses suggest that the SnRK1α1 T175 autophosphorylation occurs *in trans* and requires GRIK activity for initiation since T175 phosphorylation is completely absent in the *grik1-1 grik2-1* mutant. The latter also refutes the idea of alternative SnRK1α1-activating kinases in mature leaves under control conditions. Such kinases were proposed to exist due to restriction of GRIK protein accumulation to young, developing tissues or infected mature leaves, while SnRK1α1 T175 phosphorylation is detected in the whole plant (Kong and Hanley-Bowdoin, 2002; Shen and Hanley-Bowdoin, 2006). Possibly, SnRK1α1 is T-loop activated by GRIK proteins during early development and in rapidly growing and meristematic tissues, after which autophosphorylation maintains a pool of catalytically competent proteins with further growth under optimal conditions.

SnRK1α1 is also subject to additional non-T175 phosphorylation. One such event can be detected in the T175A kinase-dead mutant SnRK1α1 protein and in the *grik1-1 grik2-1* genetic background, indicating that it is not the result of trans-autophosphorylation or GRIK activity. The residue(s), upstream kinase(s) and signals involved remain to be identified. Interestingly, while TOR kinase is an established downstream target of SNF1/SnRK1/AMPK, human AMPKα conversely also is a TOR target and phosphorylation by TOR is associated with decreased T-loop phosphorylation (Ling et al., 2020). We also discovered another non-T175 SnRK1α1 phosphorylation event, which was only detected in the SnRK1α1-T175A mutant protein when GRIK1 was co-expressed, implying that it is not an auto-phosphorylation event but entirely dependent on GRIK1 activity. Interestingly, it was suggested that GRIK2 can phosphorylate SnRK1α1 S176 *in vitro* and that this reduced SnRK1α1 activity (Martinez-Barajas and Coello, 2019). SnRK1α1 S176 phosphorylation does occur *in planta* (Xi et al., 2021) and the residue is also highly conserved and phosphorylated *in vivo* in fungal Snf1, where it does not appear to affect kinase activity (McCartney et al., 2016), and in animal AMPKα1/2, where protein kinase A (PKA)-mediated phosphorylation results in decreased AMPK activity due to inhibition of LKB1-mediated T-loop phosphorylation (Djouder et al., 2010; Aw et al., 2014; Ferretti et al., 2016). We found that the SnRK1α1-S176D phosphomimetic, but not the -S176A mutant protein, showed compromised activity due to significantly decreased T175 phosphorylation. However, this effect was partially rescued by GRIK1 overexpression, suggesting competitive SnRK1 T-loop phosphorylation on two neighbouring residues by two antagonistic kinases. Intriguingly, a similar mechanism was recently described for the SOS2 protein, where the phosphomimetic SOS2-T169D mutant protein showed reduced activity and the phospho-dead SOS2-T169A variant showed hyperactive auto-phosphorylation and substrate phosphorylation (Li et al., 2020). Future work will explore how exactly SnRK1α1-S176 phosphorylation status controls SnRK1 activity.

As the SnRK kinases are nucleo-cytosolic proteins, we explored the localization of their upstream GRIK kinases. We discovered that both Arabidopsis GRIK1 and GRIK2 are predominantly localized at the tonoplast and that this localization relies on S-acylation of conserved cysteine residues in their N-terminal domain. Using our protoplast-based Acyl PEG Exchange (APE) protocol, we could determine the number of S-acylation events in each GRIK protein as well as identify the S-acylated Cys residues in GRIK1 (C3, C14 and C17) and GRIK2 (C14, C17). A recent study in which the Arabidopsis S-acylome was analyzed confirms S-acylation of the GRIK2 C14 and C17 residues (Kumar et al., 2022). While the S-acylated C14 and C17 residues are highly conserved in both GRIK proteins, GRIK1 harbors an additional and unique S-acylated C3 residue, which does not appear to be involved in facilitating tonoplast localization but might have another regulatory role. Mutation of the GRIK1 C3 residue did not affect localization but increased C14 and C17 S-acylation, suggesting negative cooperativity between S-acylated sites. A recent study reported that S-acylation can also result in loss of membrane association and subsequent redirection to the nucleus (Villalta et al., 2021). Alternatively, C3 S-acylation might facilitate thioesterase recruitment to remove the lipid modification at C14 and C17 and promote release from the tonoplast. GRIK1 was shown to translocate to the nucleus during leaf development and virus infection (Kong and Hanley-Bowdoin, 2002), while GRIK2 subcellular localization was predominantly non-nuclear (Shen and Hanley-Bowdoin, 2006), which supports different regulatory mechanisms for GRIK1 and GRIK2. Moreover, during heat stress SnRK1α1 appears to be T175 phosphorylated in cytosolic stress granules (SGs) (Gutierrez-Beltran et al., 2021), implying tonoplast release and recruitment of one or both GRIK proteins to these SGs. Alternatively, a different SnRK1α1-activating kinase that locates to SGs might be specifically active during heat stress. Protein S-acylation can be either stable or dynamic (He and Linder, 2009). It will be interesting to explore whether GRIK2 is stably S-acylated, while S-acylation of GRIK1 is more dynamically regulated. In addition, further research should include investigating whether and when GRIK tonoplast release occurs.

Our analyses also focused mainly on GRIK1 because of assumed redundancy. Single *grik1* and *grik2* knock out does not produce any obvious phenotype, while double *grik1-1 grik2-1* knock out is lethal (Glab et al., 2017). However, it is likely that the GRIK proteins are not fully redundant and have some unique functions. This was suggested before by differences in yeast mutant complementation and subcellular localization (Shen and Hanley-Bowdoin, 2006). Moreover, *grik1 (+/-) grik2 (-/-)* sesquimutants produce homozygous *grik1 (-/-) grik2 (-/-)* progeny, whereas the *grik1 (-/-) grik2 (+/-)* sesquimutant fails to do so (Glab et al., 2017).

While GRIK1 C3 S-acylation reduces C14 and C17 S-acylation, C14 S-acylation appears to stimulate C17 S-acylation in both GRIK proteins (and C3 S-acylation in GRIK1), indicative of positive cooperativity between S-acylated sites (Dallavilla et al., 2016; Rodenburg et al., 2017; Ulengin-Talkish et al., 2021), probably as a result of increased proximity of mono-S-acylated protein to the PATs (Rodenburg et al., 2017). Because the PATs are integral membrane proteins, initial membrane association of the substrate is a prerequisite for S-acylation to occur. We propose that the GRIK proteins utilize the hydrophobicity of their N-terminal domain to mediate transient membrane interaction. This would allow for sampling of different intracellular membranes to locate the correct partner PAT(s), which in turn S-acylate(s) the GRIK proteins for more stable attachment to the same membrane. Several lines of evidence back up this hypothesis:

First, the GRIK1 20 N-terminal residues are essential and sufficient for tonoplast localization. This sequence does not contain any alternative lipidation motifs but holds five conserved Phe (F) residues, rendering it highly hydrophobic.

Second, mutation of two Phe residues (F15A/F18A) in close proximity of the GRIK1 C14 and C17 residues, but not the C3 residue (F6A/F8A), abolishes GRIK1 S-acylation almost entirely. It is possible that the hydrophobicity of the C14 and C17 residues themselves is also important and that the conserved N-terminal CFGCFG dipeptide motif needs to be intact for transient membrane interaction and subsequent S-acylation. The absence of S-acylation in the GRIK1-F15A/F18A mutant cannot be attributed to an increased pKa/decreased reactivity of the thiols of the C14 and C17 residues. Although aromatic residues, such as phenylalanine, surrounding the cysteines can lower the thiol pKa, this occurs in combination with basic residues (Bizzozero et al., 2001), not present in the 20 N-terminal GRIK1 sequence. Various studies have shown that hydrophobic residues adjacent to cysteines also enhance S-acylation of mammalian proteins (Skene and Virag, 1989; Topinka and Bredt, 1998; El-Husseini et al., 2000). Another possibility is that the double F15A/F18A mutation disrupts an S-acylation consensus motif, that is then no longer recognized by the upstream PAT. PAT substrate recognition is still somewhat elusive and no consensus motif for S-acylation has been identified (Salaun et al., 2010). There are indications that the flanking regions of S-acylated cysteines play a role in the latter’s recognition by the respective PAT (Philippe and Jenkins, 2019; Chen et al., 2021), but a stochastic model in which S-acylation is determined by the accessibility of the cysteine to the active site of the PAT enzyme rather than by a local consensus sequence, is also proposed (Rodenburg et al., 2017).

Third, use of hydrophobicity to initiate membrane association is well established for mammalian proteins (Greaves and Chamberlain, 2006; Greaves et al., 2008, 2009; Locatelli et al., 2020) and has also been described in Arabidopsis (Konrad et al., 2014).

Fourth and finally, the Arabidopsis CBL2, CBL3 and CBL6 proteins are targeted to the tonoplast via S-acylation without alternative membrane targeting signals, similar to the GRIK proteins (Batistic et al., 2010, 2012; Zhang et al., 2016). The 22 N-terminal amino acid residues of these CBL proteins are highly conserved and were shown to mediate tonoplast targeting (Batistic et al., 2010, 2012; Zhang et al., 2016). They also include five conserved hydrophobic residues in proximity to the S-acylated Cys residues, albeit not as the GRIK CFG motif.

In conclusion and based on the results described above, we believe that this type of N-terminal hydrophobic domain represents a general feature for initial membrane association of soluble plant proteins lacking alternative signals such as transmembrane domains, prenylation, or N-myristoylation motifs. Whether these domains are self-sufficient or require chaperone proteins remains to be investigated.

Our study further suggests that the GRIK proteins localize to the tonoplast independently of a secretory pathway. Treatment with BFA or co-expression of the SAR1-H74L mutant protein, inhibitors of respectively COPI and COPII vesicles (Takeuchi et al., 2000; Pedrazzini et al., 2013), had no effect on GRIK1 S-acylation nor tonoplast localization. Moreover, inhibition of S-acylation by 2-BP resulted in GRIK1 redistribution to the nucleus and the cytoplasm, rather than to the ER, Golgi bodies or TGN. Hence, the GRIK proteins do not reach the tonoplast via sequential trafficking using compartments of the secretory pathway. The tonoplast-localized CBL proteins utilize a similar direct pathway (Batistic et al., 2012), suggesting that this might be a common route for peripheral tonoplast proteins that are lipid-modified by S-acylation only. This leads us to propose that the GRIK proteins undergo local S-acylation at their target membrane, the tonoplast. This implies that the PAT(s) responsible for GRIK S-acylation are located at the tonoplast. PAT10 and PAT11 are the sole tonoplast-localized PATs in Arabidopsis (Batistic, 2012) and hence were considered as candidates. We first explored the possibility of a dominant-negative effect of catalytically inactive PAT enzymes as observed upon overexpression in mammalian cells, likely caused by sequestration of the active endogenous PATs through oligomerization (Lai and Linder, 2013). However, the Arabidopsis CBL2 protein, that is uniquely targeted by PAT10, was still fully S-acylated with co-expression of the inactive PAT10-C192A mutant protein in leaf mesophyll protoplasts, suggesting differences between mammalian and plant PATs, despite their overall conserved function. We then turned to mutant analysis. Both GRIK proteins were still fully S-acylated in the *pat10-1* knock out background, indicating that either PAT10 is not the enzyme catalyzing GRIK S-acylation or that there exists functional redundancy between PAT10 and PAT11, and/or other PATs. Similarly, no effects were observed in two *pat11* T-DNA insertion lines, although they only show a partial reduction in gene expression, and it is unclear what the effect is at the protein level. Single Arabidopsis PAT knock out mutants often show pleiotropic phenotypes and proteomics confirm that individual PATs have many different substrates (Hemsley et al., 2013; Hemsley, 2020). Conversely, S-acylation of specific proteins can be mediated by different PATs (Gao et al., 2022; Lai et al., 2015). Thus, further investigation will require generation of higher order genuine KO mutants, somewhat complicated by the sterility of homozygous *pat10* mutants (Zhou et al., 2013).

Finally, we explored the physiological relevance of GRIK tonoplast localization, first focusing on the SOS2 pathway and salt stress response. We found that the *grik1-1 grik2-1* mutant complemented with the non-S-acylatable GRIK1-C3A/C14A/C17A (GRIK1-3C) mutant protein is hypersensitive to salinity with low germination efficiency, compromised root elongation, and significant cotyledon bleaching when grown on medium with high NaCl concentrations. Consistently, induction of the salt-responsive genes *RD22*, *RD29a* and *P5CS1* was significantly affected. We propose that the deficient salt stress response results from decreased SOS2 T168 phosphorylation and activation. While S-acylation and tonoplast localization are not required for GRIK1 activity, its ability of phosphorylating and activating SOS2 is apparently reduced. The expression of salt-responsive genes such as *P5CS* and *RD29a* is controlled by *ESE1* (Zhang et al., 2011), which in turn is induced by the SOS2-phosphorylated and -activated EIN3 transcription factor (Quan et al., 2017). Hence, reduced *ESE1* induction in the *grik1-1 grik2-1* mutant complemented with the non-S-acylatable GRIK1-3C is consistent with reduced SOS2 activity.

As both SOS2 and GRIK1-3C are nucleo-cytosolic proteins, co-localization should increase the probability of interaction, particularly when using overexpression. However, our results argue against this and suggest that the specific tonoplast location, rather than random proximity of both proteins determines their efficient interaction. This is suggestive of another protein partner that mediates this interaction. A possible candidate is the Calcineurin B-like10 (CBL10) protein. This S-acylated protein was shown to recruit SOS2 to the tonoplast (Kim et al., 2007; Chai et al., 2020), similar to the GRIK proteins. *cbl10* mutants show a salt-sensitive phenotype (Kim et al., 2007) and complementation with the non-S-acylatable CBL10-C38S could not fully rescue this phenotype (Chai et al., 2020), indicating that its tonoplast localization is important, albeit not essential, to induce an effective salt stress response. Moreover, CBL10 promotes interaction of SOS2 with the putative calcium-permeable transporter ANN4 (Ma et al., 2019) and with the BIN2 (BRASSINOSTEROID INSENSITIVE 2) kinase (Li et al., 2020), highlighting its scaffolding properties. Hence, it is possible that tonoplast-localized GRIK1 requires tonoplast-localized CBL10 for efficient interaction with and phosphorylation and activation of SOS2 during salt stress.

We also investigated the effect of GRIK1 tonoplast localization on SnRK1 signaling. Surprisingly, the *grik1-1 grik2-1* mutant complemented with non-S-acylatable GRIK1-3C showed a mild increase, rather than decrease in SnRK1 target gene induction. We speculate that, in contrast to SOS2 regulation, the nucleo-cytosolic colocalization of the GRIK1-3C mutant protein could be responsible for less restricted interaction with and hence increased phosphorylation and T-loop activation of SnRK1α1. If so, the relevance of tonoplast-associated SnRK1α1 T175 phosphorylation and spatial separation of SnRK1 activation from activity might be to prevent SnRK1 hyper-activity. Alternatively or in addition, vacuole or tonoplast-associated or -derived signals might be important for proper SnRK1 activity regulation and/or SnRK1 has important tonoplast-localized targets. In mammalian cells, the SnRK1 ortholog AMPK is recruited to the lysosome, the equivalent of the plant vacuole, where it is T-loop phosphorylated and activated by the upstream kinase LKB1 when glucose levels are low via an AMP/ADP independent mechanism (Zhang et al., 2014, 2017). With falling glucose levels, the glycolytic enzyme aldolase becomes devoid of its substrate FBP (fructose-1,6-bisphosphate) and acts as a glucose sensor, transmitting the low-glucose signal by inducing a conformational change in the associated V-ATPase-Ragulator protein complex, which in turn allows binding of AMPK and its upstream kinase LKB1 (Zhang et al., 2014, 2017; Lin and Hardie, 2018). Although the location of AMPK activation (the lysosome) is equivalent to that of SnRK1 activation (the vacuole), there are some significant differences. First, the Ragulator protein complex is not conserved in plants (Lin and Hardie, 2018). Second, a prerequisite for recruitment of AMPK to the lysosome is the N-myristoylation of the regulatory AMPKβ subunits (Oakhill et al., 2010), while in Arabidopsis the catalytic subunit SnRK1α1 appears sufficient for tonoplast-localized interaction with GRIK1. Third, lysosomal AMPK recruitment only occurs in low glucose conditions, whereas SnRK1α1 appears to be immediately phosphorylated after protein synthesis by the GRIK proteins also under optimal conditions. Despite these obvious differences, the existence of a similar glucose-sensing mechanism (at the tonoplast) in plants cannot be excluded.

In mammalian cells, different compartmentalized AMPK pools are activated depending on the severity of the nutrient or energy stress (Zong et al., 2019). While lysosomal AMPK appears to be activated by LKB1 during low glucose stress independent of nucleotide charge, a mitochondrial pool is activated by the alternative upstream kinase CaMKK2 during severe stress conditions associated with increased relative AMP levels (Schmitt et al., 2022). Our analyses indicate that there are no alternative SnRK1α1 upstream kinases for the GRIK proteins in control conditions in Arabidopsis, but we cannot exclude stress-dependent GRIK relocalization or activity of alternative SnRK1-activating kinases recruiting SnRK1 to other organelles or locations in the cell (Gutierrez-Beltran et al., 2021).

In conclusion, we show that the GRIK proteins are the major SnRK1 and SOS2 upstream kinases in Arabidopsis and that they activate both SnRK1α1 and SOS2 at the tonoplast, adding GRIK subcellular localization by N-terminal S-acylation as another level of complexity and regulation in Arabidopsis energy and salt stress signaling.

## Materials and Methods

### Plants and plant growth

GABI_713C09 (*grik1-1*), SALK_015230 (*grik2-1*), SALK_024964 (*pat10-1*), SALK_022544 (*pat11-1*), SALK_026092 (*pat11-2*) T-DNA lines were obtained from the Nothingham Arabidopsis Stock Center (NASC). Homozygous plants were selected on 1/2 MS medium including vitamins (cat. no. M0222; Duchefa Biochemie) with kanamycin and genotyped by PCR (genotyping primers in **Supplemental Table 1**). Transgenic plants (overexpressing WT or mutant GRIK1) were generated by Agrobacterium (GV3101)-mediated transformation using flower dipping as described by Zhang et al. (2006) with minor modifications and BASTA (glufosinate ammonium) selection.

Before sowing, Arabidopsis seeds were vapour-sterilized using chlorine gas (3 mL 37% HCL in 100 mL 10% bleach) for 3, 6 or 9 h for growth in soil, on solid or in liquid medium, respectively. After sterilization, seeds were stratified in 1 mL (sterile) Milli-Q for 3 days at 4°C in the dark. Unless stated otherwise, plants were grown in soil under a 12 h light/12 h dark diurnal cycle at 21°C with 75 mE cool white fluorescent light (cat. no. F17T8/TL741/ALTO; Philips) for 4 weeks.

For seedling sugar starvation assays, 10 to 15 seeds were germinated in 1 mL 1/2 MS supplemented with 1% sucrose in individual wells of a 6-well plate, with each well representing a biological replicate. Plates were incubated under continuous light (65 μE) at 21°C for 6 days. On the 6th day, medium was refreshed, and seedlings were incubated for an additional 24 h. After incubation, seedlings were starved by replacing the medium with 1 mL 1/2 MS without sucrose and harvested by flash-freezing in liquid nitrogen at different time points after sucrose removal for RT-qPCR analysis.

To assess salt stress tolerance, different assays were performed. For the germination efficiency assays, seeds were germinated on solid 1/2 MS supplemented with 1% sucrose without or with 200 mM sorbitol (osmotic control), 100 mM NaCl or 200 mM NaCl under long day (LD) conditions (16 h light/8 h dark photoperiod) for 10 days after which germination efficiency was determined. Successful germination was defined as visible radicle protrusion. For the cotyledon bleaching assays, seeds were germinated on vertical plates containing 1/2 MS supplemented with 1% sucrose under LD conditions. After three days, seedlings were transferred to fresh growth medium supplemented with 1 % sucrose without (control) or with 150 mM NaCl. Seedlings were scored four days after transfer for cotyledon greening. Green cotyledons were defined as not fully bleached. For the root bending assays, seeds were germinated on vertical plates containing 1/2 MS supplemented with 1% sucrose under LD conditions. After three days, seedlings were transferred to fresh growth medium supplemented with 1% sucrose without (control) or with 100 mM NaCl. Plates were then placed upside down for root bending and seedlings were grown under LD conditions for an additional three days, after which root elongation was measured using the ImageJ software.

For qRT-PCR analysis of salt-responsive genes, 10-15 seedlings were grown in 1 mL liquid 1/2 MS supplemented with 1% sucrose in 6-well cell culture plates under LD conditions. After 7 days, seedlings were transferred to fresh liquid 1/2 MS supplemented with 1% sucrose without or with 300 mM NaCl for an additional 3 h. Seedlings were harvested by flash-freezing in liquid nitrogen and stored until RNA isolation.

### Plasmid construction and mutagenesis

After PCR amplification (cloning primers in **Supplemental Table 1**) from *Arabidopsis thaliana* ecotype Columbia-0 (Col-0) copy DNA (cDNA), full-length coding sequences (CDS) without the STOP codon were inserted in pUC18-based expression vectors with HA-, FLAG-, GFP-or split YFP-tag for transient expression in Arabidopsis leaf mesophyll protoplasts or a pCB302-based binary vector for Agrobacterium-mediated transformation using restriction enzymes. Both vectors contain the 35SC4PPDK promoter (35S enhancer and maize C4PPDK basal promoter) and nopaline synthase (NOS) terminator. CDS site-directed mutagenesis of was done by PCR using Phusion polymerase (Thermo Fisher) and complementary primers containing the required modification(s) flanked by 15 matching nucleotides on both sides (**Supplemental Table 1**).

### Protoplast isolation and transfection

Arabidopsis leaf mesophyll protoplast isolation and transfection were performed essentially according to Yoo et al. (2007). Depending on the type of experiment, different volumes of protoplasts (50 μL for LUC assays, 100 μL for immunoblotting, 200 μL for confocal microscopy and 1 μL for RNA isolation; 2x 10^5^ protoplasts/mL) were transfected with proportionally different volumes of CsCl gradient-purified plasmid DNA and reagents. Transfection of protoplasts with the plasmid DNA construct(s) of interest was performed by adding 1 volume PEG-Ca^2+^ solution (40% [w/v] PEG 4000 (Sigma Aldrich), 0.2 M D-mannitol, 100 mM CaCl2) to the protoplast-DNA mixture. After gentle tapping to ensure homogeneity and 5 min incubation at RT, transfection was stopped by addition of 2 volumes of W5 solution (2 mM MES pH 5.7, 154 mM NaCl, 125 mM CaCl2, 5 mM KCl), followed by centrifugation for 1 min (100 g, RT). The protoplast pellet was resuspended in a small volume of WI incubation buffer (4 mM MES pH 5.7, 0.5 M mannitol, 20 mM KCl) and incubated in WI buffer in 5% calf serum pre-coated cell culture plates. After 6 h incubation, cells were harvested by centrifugation (1 min, 200 g, RT) and stored at − 80°C. Protoplasts for confocal fluorescence microscopy were incubated for 14 to 16 h and immediately analyzed after harvest.

### Luciferase-based (LUC) promoter activity assays

*SnRK1* activity was investigated using a luciferase (LUC) reporter constructs with the coding sequence of firefly luciferase (LUC) under control of the *DIN6/ASPARAGINE SYNTHASE1 (At3g47340)* promoter (Baena-Gonzalez et al., 2007). A pr*UBQ10::GUS* reporter construct containing the constitutively active *UBIQUITIN10 (UBQ10)* promoter fused to the coding sequence of β-glucuronidase (GUS) was used as an internal control to compensate for pipetting errors and differences in transfection efficiency. 50 μL 1x Cell Culture Lysis Reagent (25 mM Tris/phosphate pH 7.8, 2 mM DTT, 2 mM trans-1,2-diaminochyclohexane-N,N,N’,N’-tetraacetic acid (DACTAA), 10% [v/v] glycerol, 1% [v/v] Triton X-100) (Promega) was added to the harvested protoplasts for lysis. 25 μL lysate was transferred into a plastic tube that was placed into a luminometer (Lumat LB 9507, Berthold Technologies), which automatically adds 100 μL of 1x Luciferase Assay Reagent (Promega) to the sample and subsequently measures the luminescence ((background: 0.5 s, max. 150 relative light units (RLU)/s, delay: 2 s, measurement: 10 s). For the GUS assay, 5 μL lysate was mixed with 45 μL MUG (methylumbelliferyl b-D-glucuronide) solution (10 mM Tris/HCl pH 8, 2 mM MgCl2, 1 mM MUG (Sigma Aldrich)) in a 96-well plate. The plate was incubated at 37 C for 1 h, after which 220 μL 0.2 M Na2CO3 was added to stop the reaction. GUS activity was measured (excitation wavelength MU: 365 nm, emission wavelength MU: 455 nm) using a fluorometer (Glomax-Multi+ Detection System with InstinctR Software, Promega). The promoter activity was determined by calculating the relative LUC/GUS ratio.

### Immunoblotting

For confirmation of protein expression, standard SDS-PAGE followed by wet blotting and visualization with the appropriate antibodies was used. 20 μL loading buffer (62.5 mM Tris/HCL pH 6.8, 6Murea, 10%[v/v] glycerol, 2%[v/v] SDS, 5%[v/v] β-mercaptoethanol (BME), and 1% [v/v] bromophenol blue (BPB)) was added directly to the protoplast pellet. Samples were then briefly vortexed and heated for 5 min at 95 C prior to loading on a manually casted 1.5 mm discontinuous SDS-PAGE gel consisting of a 7.5% or 10% running gel and a 5% stacking gel (Table 6.4). As a protein size reference, the SeeBlue Plus2 Prestained Protein Standard (Thermo Fisher) was used. Proteins were separated in Tris-Gly running buffer (25 mM Tris/HCl, 192 mM glycine, 0.5% [w/v] SDS) at 60 V for 15 min, followed by 110 V for 15 min and 160 V until the BPB band reached the bottom of the running gel. Proteins were then transferred to a methanol (MeOH)-activated polyvinylidene fluoride (PVDF) membrane (Immobilon-P; Millipore) using a wet tank electro-blotting apparatus (Mini Trans-Blot; Bio-Rad) in Tris-Gly transfer buffer (25 mM Tris/HCl, 192 mM glycine, 0.5% [w/v] SDS, 10% [v/v] MeOH) at 300 mA for 2 h. To prevent overheating, cooling units were placed into the tank, and refreshed after 1 h. After wet blotting, the membrane was first incubated for 1 h in 50 mL 5% [w/v] semi-skimmed milk in TBS-T (150 mM NaCl, 20 mM Tris/HCl pH 7.6, 0.05%[v/v] Tween 20 (Merck) at room temperature (RT), followed by 2 h incubation with (primary) antibody in 10 mL 1% milk solution at RT [horseradish peroxidase (HRP)-conjugated anti-HA antibody, 1/1000 (Roche); HRP-conjugated anti-FLAG antibody, 1/1000 (Sigma-Aldrich)]. The membrane was washed three times with TBS-T, incubated in Pierce SuperSignal West Pico PLUS Chemiluminescent Working Solution (Thermo Fisher Scientific) for 5 min, exposed to light-sensitive film (CL-Xposure; Thermo Fisher) for a few seconds to several minutes, and developed manually. After development, the membrane was stained with Coomassie blue solution [0.1% [w/v] Coomassie R-250 (Thermo Fisher), 40% [v/v] ethanol (EtOH), 10% [v/v] acetic acid)] for total protein visualization. Ribulose bisphosphate carboxylase large chain (RBCL) staining of the blot with Coomassie Brilliant Blue R-250 was used as a protein loading control.

### Phos-tag mobility shift assays

For the visualization of protein phosphovariants, Phos-tag mobility shift assays were performed. A Phos-tag SDS-PAGE gel consisted of a 7.5% or 10% running gel supplemented with 25 or 50 μM Phos-tag (AAL-107; Fujifilm Wako Chemicals) reagent and 50 or 100 μM MnCl_2_ respectively, and a 5% stacking gel. Electrophoresis was performed in Tris-Gly running buffer, first at constant voltage (60 V for 15 min, 110 V for 15 min), then at constant current of 30 mA/gel for at least 2 h. After protein separation, gels were incubated 2x 20 min with Tris-Gly transfer buffer containing 10 mM EDTA to eliminate Mn^2+^, followed by one wash for 10 min in Tris-Gly transfer buffer without EDTA. Proteins were then transferred to a PVDF membrane using wet blotting and visualized with chemiluminescence as described for immunoblotting above.

### Protein stability assay

For the protein stability assay, 200 μL (4 x 10^4^) protoplasts were transfected with 20 μL (40 μg) plasmid DNA. After transfection, the sample was split in four (one sample for each time point) and incubated. After 4 h cycloheximide (CHX) was added to a final concentration of 20 μM to all samples and cells were harvested at different time points by centrifugation at 100 g for 1 min and subjected to immunoblotting as described for above.

### Confocal fluorescence microscopy

To determine subcellular localization of GFP-and mRFP-tagged proteins, 200 μL (4 x 10^4^) protoplasts were transfected with 20 μL (40 μg) plasmid DNA and incubated for 14 to 16 h. Similarly, for BiFC assays, 200 μL (4 x 10^4^) protoplasts were transfected with 10 μL (20 μg) of each BiFC(split YFP) plasmid and incubated for 14-16 h. GFP, YFP and mRFP fluorescence was visualized using confocal laser scanning microscopy (FV1000; Olympus) with a 40x (1.3 oil) objective.

### Acyl-PEG Exchange (APE) assay

This optimized protocol is based on the protocol developed by Percher et al. (2018) with modifications for use with the Arabidopsis protoplast experimental setup. 200 μL (4 x 10^4^) protoplasts were transfected with a total of 20 μL (40 μg) CsCl-purified DNA and incubated for 6 h in 12-well plates with 500 μL WI buffer per well. After incubation, protoplasts were harvested and lysed with 100 μL freshly prepared lysis buffer [1x TEA (5 mM triethanolamine, 150 mM NaCl, pH 7.3; a 10x buffer can be stored up to 1 year at RT], 4% SDS, 1x protease inhibitor (Roche; EDTA-free), 5 mM PMSF (phenylmethanesulfonylfluoride; Sigma), 5 mM EDTA) by gently pipetting up and down. For reducing any disulfide bridges, 92.5 μL lysate was transferred to a fresh 1.5 mL tube to which 5 μL of 200 mM neutralized (pH 7.3) TCEP (tris(2-carboxyethyl)phosphine, Thermo-Fisher) was added. The mixture was incubated at RT for 45 min with gentle rotation. For the next step, i.e. cysteine free thiol capping reaction, 2.5 μL NEM (N-ethylmaleimide, Sigma-Aldrich) of a 2 M stock (in EtOH) was added, followed by incubation for 2.5h at RT and under gentle rotation. To terminate the reaction after 2.5h, proteins were purified using MeOH-chloroform-H_2_O (4:1.5:3, relative to sample volume) precipitation by adding in the following order (all pre-chilled): 400 μL MeOH, 150 μL chloroform and 300 μL Milli-Q water. Mixtures were inverted several times and then centrifuged for 10 min at 20 000 g at 4°C. After centrifugation, a protein pellet should be visible between two separate phases. The aqueous upper layer was removed by pipetting without disturbing the protein pellet, after which 1 mL MeOH was added. Samples were inverted a few times to dislodge the pellet, followed by centrifugation at 20 000 g for 10 min at 4°C. The supernatant was decanted, the pellet was washed with 500 μL MeOH and dried for 10 min at RT. For hydroxylamine (NH_2_OH) cleavage, the protein pellet was resuspended in 60 μL resuspension buffer by gently pipetting up and down (1x TEA, 4% SDS, 4 mM EDTA). To 30 μL of the resuspended protein pellet, 90 μL of a 1 M neutralized (pH 7.3) NH_2_OH (dissolved in 1x TEA buffer containing 0.2% Triton X-100) was added. Samples were incubated for 1h at RT under gentle rotation. After incubation, NH_2_OH was removed using the previously described MeOH-chloroform-H_2_O precipitation. For the mPEG-mal alkylation, the protein pellet was resuspended in 30 μL resuspension buffer by gently pipetting up and down. 90 μL of 1x TEA containing 0.2% Triton X-100 and 1.33 mM mPEG-mal (5 kDa mPEG-maleimide, Sigma-Aldrich) was added to the samples, which were then incubated for 2h at RT with gentle rotation. After incubation, a final MeOH-chloroform-H_2_O precipitation was performed, after which 50 μL of loading buffer was added. Samples were then heated for 5 min at 95°C and 15 μL sample was loaded onto a 10% SDS-PAGE gel for subsequent immunoblotting.

### RT-qPCR

RT-qPCR was performed using the PowerUP SYBR Green Master Mix kit (Applied Biosystems) according to the manufacturer’s manual. Briefly, the 10 μL reaction mixture contained 5 μL PowerUP SYBR Green buffer, 0.2 μL 10 μM forward and reverse primer (RT-qPCR primers in **Supplemental table 1**), 2 μL 5 ng μL^−1^ cDNA template and 2.6 μL Milli-Q. The RT-qPCR thermal cycling conditions were 2 minutes at 50°C, 2 minutes at 95°C and 40 cycles of 3 seconds at 95°C and 30 seconds at 60°C. Expression levels were normalized to *UBQ10* expression, which proved stable in the used tissues and stress conditions (Baena-Gonzalez et al., 2007).

