## Supplementary Material for "S-acylation and tonoplast localization of the Geminivirus Rep-Interacting Kinase/SnRK1-Activating Kinase (GRIK/SnAK) proteins differentially regulate salt and energy stress responses in Arabidopsis"

### Supplemental Figures

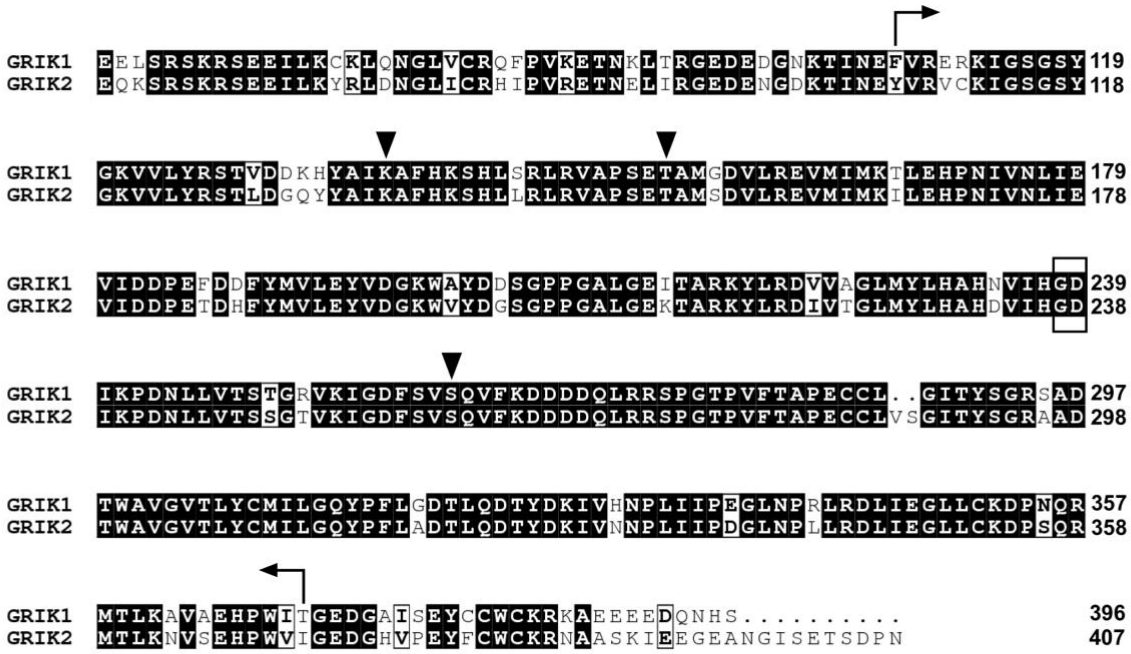


***Supplemental Figure 1: Amino acid sequence alignment of the Arabidopsis GRIK proteins.*** *The catalytic kinase domain is indicated with arrows, while the catalytic aspartate (D239/D238 in GRIK1/2 respectively) and the preceding glycine (non-arginine) residue are framed. Important residues are indicated with black arrowheads: the T154/T153 autophosphorylation site (Crozet et al., 2010), the K137/K136 involved in phosphotransfer (Shen and Hanley-Bowdoin, 2006), and the S261/260 which is phosphorylated by SnRK1 (Crozet et al., 2010). Identical residues are boxed; residues with similar physico-chemical properties are in bold. Alignment was performed with T-Coffee (Notredame et al., 2000) and visualization using ESPript (Robert and Gouet, 2004).*

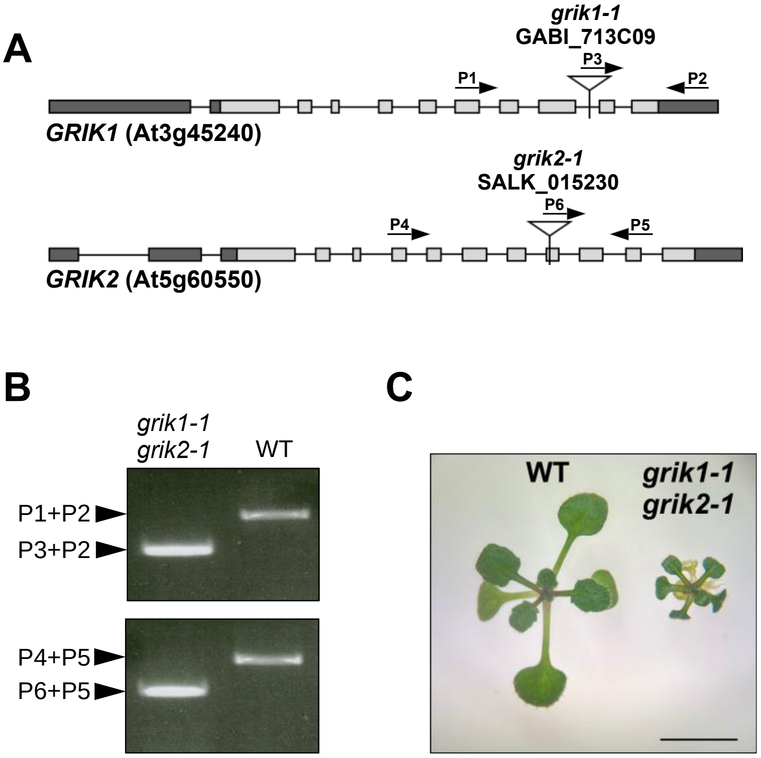


***Supplemental Figure 2: Selecting double homozygous grik1-1 grik2-1 mutants for leaf mesophyll protoplast isolation.*** *(A) Schematic representation of the two GRIK genes and the location of the T-DNA insertion) in grik1-1 and grik2-1. Light grey boxes represent exons, dark grey boxes indicate 5' UTR and 3' UTR and lines indicate introns. Triangles denote the location of the T-DNA insertion. (B) Genotyping of the grik1-1 grik2-1 double mutant. The locations of gene-specific primers are indicated in A. WT, wild type. (C) Progeny of the grik1-1 (+/-) grik2-1 (-/-) sesquimutant and WT were grown on 1/2 MS supplemented with 3% glucose for 4 weeks in a growth chamber with 12 h light/12 h dark photoperiod. grik1-1 (-/-) grik2-1 (-/-) mutant seedlings are identified by their smaller size compared to WT. Scale bar represents 0.5 cm.*

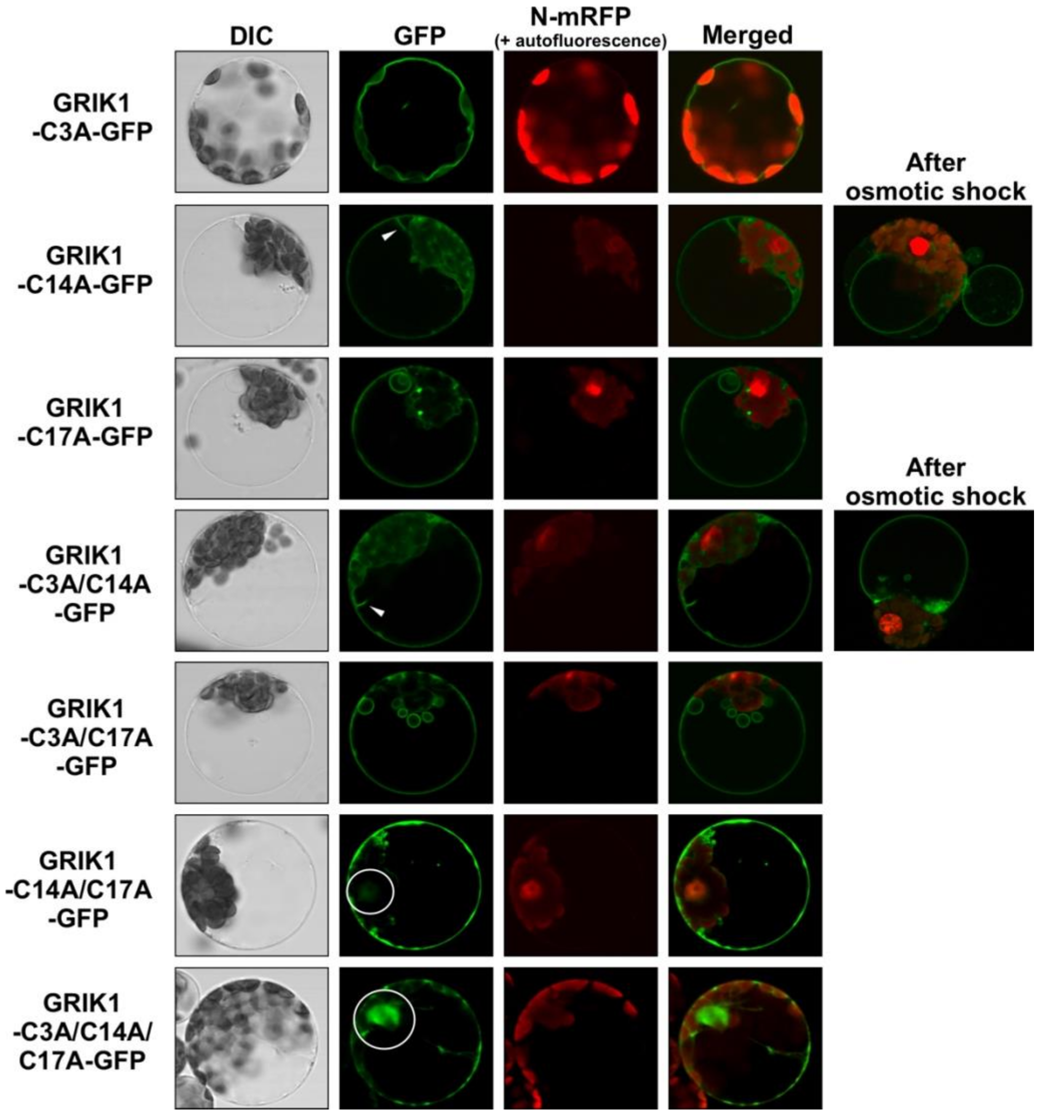


***Supplemental Figure 3: GRIK1 tonoplast targeting requires the C14 and C17 residues.*** *Confocal fluorescence microscopy of the subcellular localization of transiently expressed GFP-tagged C3A, C14A and C17A single, double and triple GRIK1 mutant proteins in leaf mesophyll protoplasts. Intact vacuoles were released using mild osmotic shock to highlight tonoplast localization. White arrowheads indicate tonoplast strands, white circles indicate the nucleus. mRFP-tagged SCF30 (Splice Factor 30) was used as a nuclear reporter (N-mRFP).*

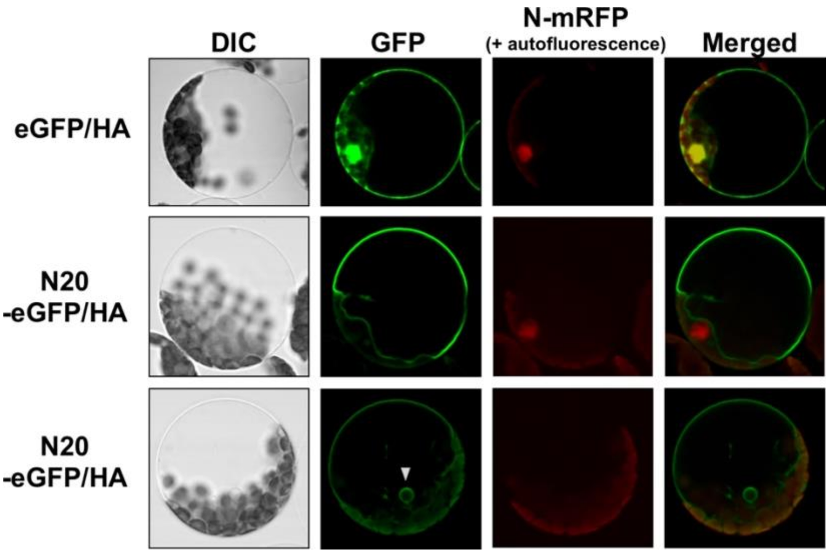


***Supplemental Figure 4: The 20 N-terminal amino acids of the GRIK1 protein are sufficient for tonoplast targeting.*** *Confocal fluorescence microscopy of subcellular localization of transiently expressed WT eGFP/HA and N20-eGFP/HA fusion proteins in leaf mesophyll protoplasts. The white arrowhead indicates a tonoplast bulb.*

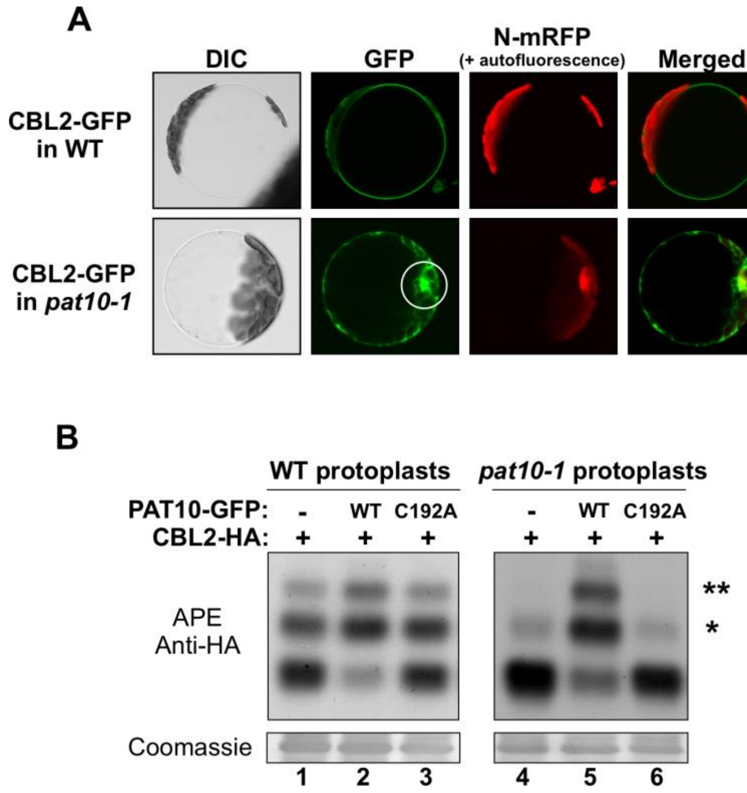


***Supplemental Figure 5: Validation of the protoplast-based APE protocol by PAT10-mediated CBL2 S-acylation.*** *(A) Confocal fluorescence microscopy of transiently expressed GFP-tagged CBL2 localization in WT and in pat10-1 leaf mesophyll protoplasts. The white circle indicates the nucleus. mRFP-tagged SCF30 (Splice Factor 30) was used as a nuclear reporter (N-mRFP). (B) APE immunoblot assay of WT and pat10-1 mutant leaf mesophyll protoplasts transiently expressing HA-tagged CBL2 without or with GFP-tagged WT PAT10 or mutant inactive PAT10-C192A proteins. Immunoblot analyses were performed using anti-HA antibodies. Coomassie blue staining of the large subunit of Rubisco serves as loading control. The number of PEGylations corresponds to the number of S-acylation events represented by the asterisks.*

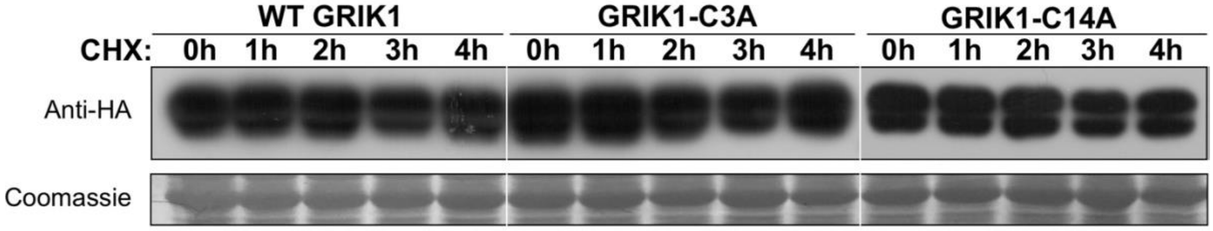


***Supplemental Figure 6: GRIK1 C3A or C14A mutation does not affect protein stability.*** *Leaf mesophyll protoplasts were transfected with HA-tagged WT GRIK1, GRIK1-C3A or GRIK1-C14A constructs and incubated for 4 h after which cycloheximide (CHX) was added to a final concentration of 20 µM in the incubation medium. Protoplasts were harvested at different time points after addition of CHX. Immunoblot analyses were performed using anti-HA antibodies. Coomassie blue staining of the large subunit of Rubisco served as loading control.*

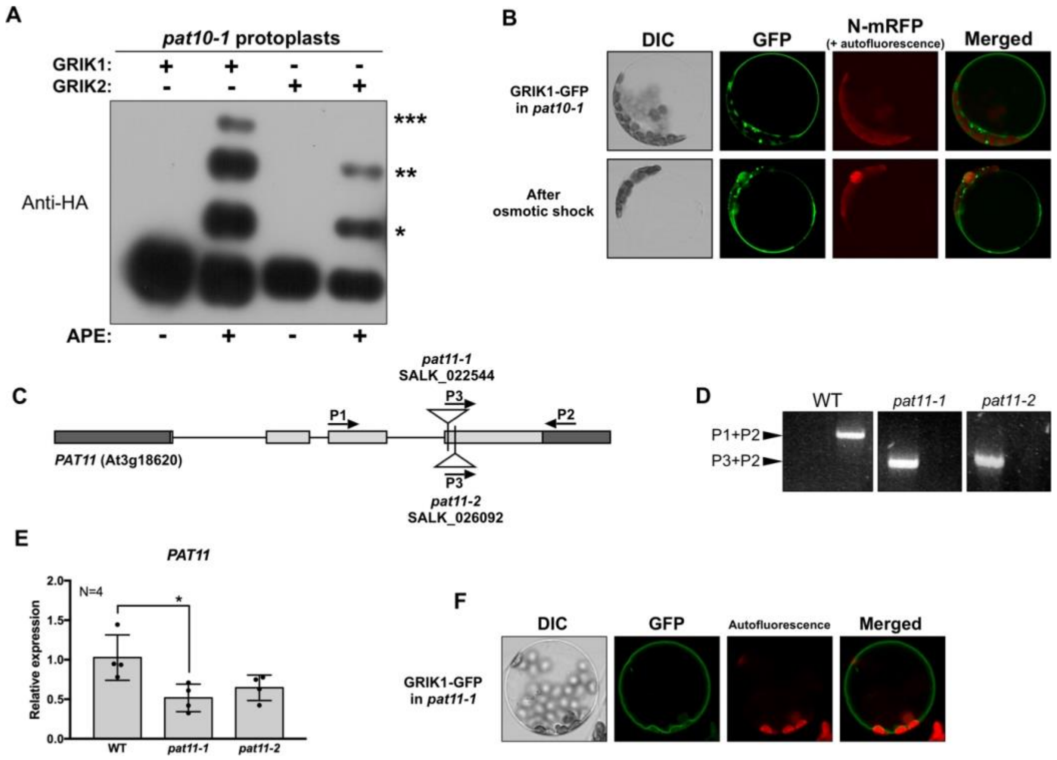


***Supplemental Figure 7: GRIK S-acylation and tonoplast localization in pat10 and pat11 mutants***

*(A) Immunoblot of the APE assay with pat10-1 leaf mesophyll protoplasts transiently expressing HA-tagged WT GRIK1 or GRIK2. The number of PEGylations corresponds to the number of S-acylation events and is indicated by asterisks. Proteins were visualized using anti-HA antibodies. (B) Confocal fluorescence microscopy of subcellular localization of transiently expressed GFP-tagged GRIK1 in pat10-1 leaf mesophyll protoplasts. mRFP-tagged SCF30 (Splice Factor 30) was used as a nuclear reporter (N-mRFP). Intact vacuoles were released using a mild osmotic shock to highlight tonoplast localization. (C) Schematic representation of the PAT11 gene and location of the T-DNA insertions in the pat11-1 and pat11-2 mutant lines. Light grey boxes represent exons, dark grey boxes indicate 5' UTR and 3' UTR and lines indicate introns. Triangles denote the T-DNA insertion location. (D) Genotyping of the pat11-1 and pat11-2 T-DNA insertion lines. The locations of gene-specific primers are indicated in C. (E) qRT-PCR analysis of PAT11 expression in rosette leaves of 4-week-old WT, pat11-1 and pat11-2 plants. Values are averages with standard deviations. N=4 biological repeats (2 leaves pooled per repeat). Statistical analyses were performed with Graphpad Prism 7. One-way ANOVA, *P<0.1. (F) Confocal fluorescence microscopy of subcellular localization transiently expressed GFP-tagged GRIK1 in pat11-1 leaf mesophyll protoplasts. mRFP-tagged SCF30 (Splice Factor 30) was used as a nuclear reporter (N-mRFP). Intact vacuoles were released using a mild osmotic shock to highlight tonoplast localization.*

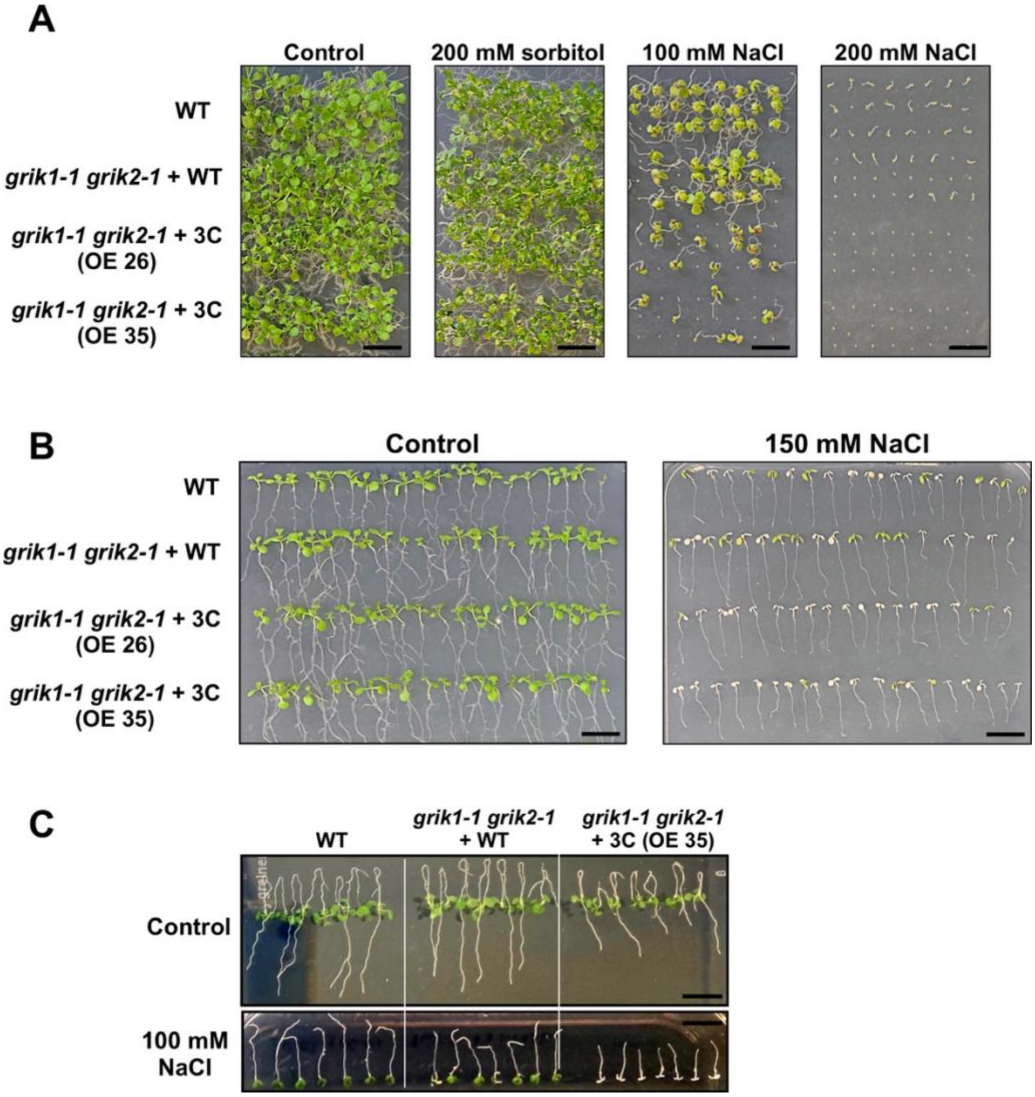


***Supplemental Figure 8: Loss of GRIK1 S-acylation and tonoplast localization increases Arabidopsis salt stress sensitivity.*** *Effect of salt stress on germination efficiency (A), cotyledon greening (B) and root elongation (C) in WT and grik1-1 grik2-1 mutant lines complemented with WT GRIK1 or non-acylatable mutant GRIK1-3C as described and quantified in Figure 7. NaCl and sorbitol (osmotic control) concentrations are indicated. Scale bars represent 0.5 cm.*

### Supplemental Tables

***Supplemental Table 1: PCR primers used in this study***

*Genotyping*

| | | |
|---|---|---|
| *GABI_713C09-LP* | *ATACATGACCGACAGACCGAG* | *GRIK1/At3g45240* |
| *GABI_713C09-RP* | *TCATGCTCATGTATGCTCTGC* | *GRIK1/At3g45240* |
| *GABI-LB* | *ATATTGACCATCATACTCATTGC* | *T-DNA GABI line* |
| *SALK_015230-LP* | *TTTCCAGGCATTTCACAAGTC* | *GRIK2/At5g60550* |
| *SALK_015230-RP* | *ATCCATCCGGGATTATCAGAG* | *GRIK2/At5g60550* |
| *SALK_022544-LP* | *TGCATATCCCGTCTTGTTTTC* | *PAT11/At3g18620* |
| *SALK_022544-RP* | *GAATTGACCACAACAAGCTCC* | *PAT11/At3g18620* |
| *SALK_024964-LP* | *ATTTGACATGCGGCTATTGTC* | *PAT10/At3g51390* |
| *SALK_024964-RP* | *TATCGATGCAGGTGTAGGGTC* | *PAT10/At3g51390* |
| *SALK_026092-LP* | *TGCATATCCCGTCTTGTTTTC* | *PAT11/At3g18620* |
| *SALK_026092-RP* | *GAATTGACCACAACAAGCTCC* | *PAT11/At3g18620* |
| *SALK-LB* | *ATTTTGCCGATTTCGGAAC* | *T-DNA SALK line* |

*Cloning*

| | | |
|---|---|---|
| *aMAN1-A* | *CGGGATCCATGGCGAGAAGTAGATCGATTA* | *aMAN1b/At1g51590* |
| *aMAN1-B* | *AAGGCCTAACGTTAATCTGATGACCAAACT* | *aMAN1b/At1g51590* |
| *CBL2-A* | *CGGGATCCATGTCGCAGTGCGTTGACGG* | *CBL2/At5g55990* |
| *CBL2-B* | *AAGGCCTGGTATCTTCAACCTGAGAATGG* | *CBL2/At5g55990* |
| *GRIK1-A* | *CGGGATCCATGTTTGTGATAGTTTTGCATTTG* | *GRIK1/At3g45240* |
| *GRIK1-B* | *AAGGCCTGCTATGGTTTTGATCTTCTTCTTC* | *GRIK1/At3g45240* |
| *PAT10-A* | *CGGGATCCATGGGCGTTTGTTGCCCTTTC* | *PAT10/At3g51390* |
| *PAT10-B* | *GCGATATCGCAGCAGCGACATTTCAACATA* | *PAT10/At3g51390* |
| *PAT11-A* | *CGGGATCCATGGAAGATTCTTCCCAGGGG* | *PAT11/At3g18620* |
| *PAT11-B* | *AAGGCCTTGTCTTATGTCTCTTCCTCAAGT* | *PAT11/At3g18620* |
| *SAR1B-A* | *CGGGATCCATGTTCTTGTTCGATTGGTTCTA* | *SAR1B/At1g28580* |
| *SAR1B-B* | *AAGGCCTGTTGATGTACTGAGAGAGCCA* | *SAR1B/At1g28580* |

*Mutagenesis*

| | | |
|---|---|---|
| *GRIK1-C3A-A* | *CGGGATCCATGTTTGCTGATAGTTTTGCATTTG* | *GRIK1/At3g45240* |
| *GRIK1-C14A-A* | *GCCCAAGTGATGAGTGCCTTCGGGTGTTTTGG* | *GRIK1/At3g45240* |
| *GRIK1-C14A-B* | *CCAAAACACCCGAAGGCACTCATCACTTGGGC* | *GRIK1/At3g45240* |
| *GRIK1-C17A-A* | *TGAGTTGCTTCGGGGCTTTTGGTGGCTCTG* | *GRIK1/At3g45240* |
| *GRIK1-C17A-B* | *CAGAGCCACCAAAAGCCCCGAAGCAACTCA* | *GRIK1/At3g45240* |
| *GRIK1-C14A/C17A-A* | *GCCCAAGTGATGAGTGCCTTCGGGGCTTTTGG* | *GRIK1/At3g45240* |
| *GRIK1-C14A/C17A-B* | *CCAAAAGCCCCGAAGGCACTCATCACTTGGGC* | *GRIK1/At3g45240* |
| *GRIK1-K137A-A* | *ACAAGCATTATGCTATTGCGGCTTTTCACAAGTCAC* | *GRIK1/At3g45240* |
| *GRIK1-K137A-B* | *GTGACTTGTGAAAAGCCGCAATAGCATAATGCTTGT* | *GRIK1/At3g45240* |
| *GRIK2-C14A-A* | *GCCCGTACCATCGGTGCCTTTGGATGCTTTGG* | *GRIK2/At5g60550* |
| *GRIK2-C14A-B* | *CCAAAGCATCCAAAGGCACCGATGGTACGGGC* | *GRIK2/At5g60550* |
| *GRIK2-C17A-A* | *ATCGGTTGCTTTGGAGCCTTTGGCAGTTCTGG* | *GRIK2/At5g60550* |
| *GRIK2-C17A-B* | *CCAGAACTGCCAAAGGCTCCAAAGCAACCGAT* | *GRIK2/At5g60550* |
| *GRIK2-C14A/C17A-A* | *ATCGGTGCCTTTGGAGCCTTTGGCAGTTCTGG* | *GRIK2/At5g60550* |
| *GRIK2-C14A/C17A-B* | *CCAGAACTGCCAAAGGCTCCAAAGGCACCGAT* | *GRIK2/At5g60550* |
| *PAT10-C192A-A* | *CAGTTTGATCATCACGCTGTTTGGTTAGGAAC* | *PAT10/At3g51390* |
| *PAT10-C192A-B* | *GTTCCTAACCAAACAGCGTGATGATCAAACTG* | *PAT10/At3g51390* |
| *PAT11-C178S-A* | *GACATGGATCACCATAGCCCCTTTATTGGGA* | *PAT11/At3g18620* |

| | | |
|---|---|---|
| *PAT11-C178S-B* | *TCCCAATAAAGGGGCTATGGTGATCCATGTC* | *PAT11/At3g18620* |
| *SAR1B-H74L-A* | *TTGATTTGGGTGGTCTTCAGATTGCTCGTAG* | *SAR1B/At1g28580* |
| *SAR1B-H74L-B* | *CTACGAGCAATCTGAAGACCACCCAAATCAA* | *SAR1B/At1g28580* |

*RT-qPCR*

| | | |
|---|---|---|
| *ATG8e-qRT-A* | *GCATCTTTAAGATGGACGACGATTTCGAA* | *ATG8e/At2g45170* |
| *ATG8e-qRT-B* | *ATGTGTTCTCGCCACTGTAAGTGATGTAA* | *ATG8e/At2g45170* |
| *BCAT2-qRT-A* | *TCACAAATTATGCGCCAGTT* | *BCAT2/At1g10070* |
| *BCAT2-qRT-B* | *CGAGATAAAGAACGTCTGAAAACC* | *BCAT2/At1g10070* |
| *DIN6-qRT-A* | *AACTTGTCGCCAGATCAAGG* | *DIN6/At3g47340* |
| *DIN6-qRT-B* | *GGAACACGTGCCTCTAGTCC* | *DIN6/At3g47340* |
| *ESE1-qRT-A* | *ACGTTTAACACAGCGGAAGA* | *ESE1/At3g23220* |
| *ESE1-qRT-B* | *TTGGACCGTCCTTCATCATTT* | *ESE1/At3g23220* |
| *PAT11-qRT-A* | *CATGGATCACCATTGCCCCT* | *PAT11/At3g18620* |
| *PAT11-qRT-B* | *AGCTTGTGCTAATGACGGCT* | *PAT11/At3g18620* |
| *P5CS1-qRT-A* | *AGCAGCCTGTAATGCGATGG* | *P5CS1/At2g39800* |
| *P5CS1-qRT-B* | *AAGTGACGCCTTTGGTTTGC* | *P5CS1/At2g39800* |
| *RD22-qRT-A* | *AGGGCTGTTTCCACTGAGG* | *RD22/At5g25610* |
| *RD22-qRT-B* | *CACCACAGATTTATCGTCAGACA* | *RD22/At5g25610* |
| *RD29A-qRT-A* | *GTTACTGATCCCACCAAAGAAGA* | *RD29A/At5g52310* |
| *RD29A-qRT-B* | *GGAGACTCATCAGTCACTTCCA* | *RD29A/At5g52310* |
| *UBQ10-qRT-A* | *AACTTTGGTGGTTTGTGTTTTGG* | *UBQ10/At4g05320* |
| *UBQ10-qRT-B* | *TCGACTTGTCATTAGAAAGAAAGAGATAA* | *UBQ10/At4g05320* |
